# Computationally Driven Top-Down Mass Spectrometry of Ubiquitinated Proteins

**DOI:** 10.1101/2025.07.24.666707

**Authors:** Elizaveta I. Shestoperova, Daniil G. Ivanov, Joyce Zhong, Liam Chien, Eric R. Strieter

**Author notes:** Corresponding Author: Eric R. Strieter.

## Abstract

Ubiquitination regulates numerous cellular processes through the attachment of polyubiquitin (Ub) chains that vary in linkage type, length, and branching topology. However, current mass spectrometry approaches cannot simultaneously define both the site of ubiquitination and the topology of the attached Ub chain on intact protein substrates. Here, we present the first integrated strategy that enables simultaneous determination of ubiquitin site and chain architecture using top-down mass spectrometry (TD-MS). Central to this approach is ***UbqTop***, a custom computational platform that predicts Ub chain topology from tandem MS (MS²) fragmentation data by utilizing Bayesian-like scoring algorithm. To address the challenge of analyzing complex substrates, we combine this with selective Asp-N proteolysis, which digests the substrate while preserving intact Ub chains. This enables direct, site-resolved mapping of Ub chain topology on proteins. We demonstrate the broad utility of this method on both free Ub chains and multiply ubiquitinated protein substrates, including the resolution of isomeric chains and branched architectures. Together, this work establishes a powerful new framework for proteoform-level analysis of ubiquitin signaling with unprecedented structural resolution.

## INTRODUCTION

Ubiquitination is a posttranslational modification in eukaryotic cells that regulates a wide range of biological processes ^1–4^. Through an enzymatic cascade involving E1, E2, and E3 enzymes, ubiquitin (Ub) is covalently attached to protein substrates ^5^. Ub itself can form polymeric chains through isopeptide bonds between the ε-amino group of one of its internal lysine residues (K6, K11, K27, K29, K33, K48, or K63) or its N-terminal methionine (M1), and the C-terminal carboxyl group of another Ub molecule. This results in a diverse array of Ub conjugates that vary in length, linkage type, and degree of branching ^6–10^. The specific architecture of a Ub chain is closely linked to the biological outcome of ubiquitination ^13–20^. However, the structural complexity and heterogeneity of Ub chains present a major challenge to systematically dissecting how chain type influences Ub function ^21^.

Mass spectrometry (MS)–based approaches have been instrumental in characterizing protein ubiquitination ^22^. In bottom-up proteomics, proteins are digested into small peptides, allowing identification of Ub linkage types and modification sites through detection of diGly remnants on Ub and substrate lysines ^23–26^. Middle-down proteomics offers additional insight by employing limited proteolysis to generate Ub conjugates with partial chain structures and characteristic diGly signatures that can suggest branching ^27–30^. However, both bottom-up and middle-down strategies fragment the original Ub conjugate, resulting in the loss of information about overall chain architecture, including topology, branching, and length ^21^. In the case of substrate-conjugated Ub chains, these methods also disrupt the spatial relationship between chain type and its attachment site—an important limitation, as the placement of a specific chain on a given substrate site can dictate its processing by deubiquitinases or its recognition by Ub-binding effectors, ultimately leading to distinct biological outcomes ^31–33^. Therefore, fully resolving the functional logic of ubiquitination requires an approach that preserves both the topology of the chain and its connectivity to the intact substrate.

Top-down mass spectrometry (TD-MS) holds significant potential for the analysis of ubiquitinated proteins ^21^. By preserving proteins in their intact form, TD-MS retains critical information about Ub chain length, linkage composition, topology, and connectivity to the substrate. Several pioneering studies have applied TD-MS to characterize both free and substrate-conjugated Ub chains ^34–36^. However, these efforts often rely on multistep, manual workflows that sequentially evaluate fragment coverage for each Ub subunit to infer chain identity ^37^. While informative, such analyses are labor-intensive and not scalable, limiting the broader application of TD-MS for Ub chain analysis. Moreover, overlapping fragment patterns within Ub chains reduce the effectiveness of conventional fragment-matching algorithms, which typically consider only the presence of detected ions and are often insufficient to distinguish among Ub isomers ^38^.

To address these challenges, we developed a fully automated platform for Ub chain topology analysis, implemented in the software *UbqTop*. This algorithm generates theoretical lists of MS² fragments for user-defined Ub chains or ubiquitinated substrates and compares them to experimental spectra, followed by a probability calculation using a Bayesian-inspired scoring system that accounts for both fragment presence and absence. *UbqTop* enables rapid, unbiased prediction of Ub chain architecture and was validated on a comprehensive panel of free Ub chains varying in length, linkage, and branching topology. To extend this strategy to protein substrates, we identified the protease Asp-N as uniquely suited for native digestion: it selectively cleaves substrates without disrupting Ub chains, reducing substrate mass while maintaining linkage between the substrate and intact polyUb chains. This facilitates high-quality TD-MS acquisition and enables mapping of chain topology directly to specific ubiquitination sites. This is the first approach capable of simultaneously identifying both the site of ubiquitination and the full topology of the attached chain—capabilities that cannot be achieved by bottom-up, middle-down, or existing top-down methods. By overcoming longstanding barriers in Ub proteoform analysis, our integrated experimental and computational workflow establishes a generalizable framework for high-resolution, site-specific deconvolution of ubiquitin signaling.

## RESULTS

### Analytical framework for enumeration of theoretical fragments of Ub conjugates

Automated annotation of TD-MS data for Ub conjugates requires a machine-readable representation of chain topology. Unlike linear polypeptides, which can be described using standard single-letter amino acid sequences, Ub chains exhibit non-linear, branched architectures that demand a more flexible data structure. To address this, we implemented a graph-based representation in the form of oriented labeled trees, defined as *G* = (*V*, *E*), where *V* is the set of Ub subunits (nodes) and, and *E* is the set of isopeptide bonds (edges) connecting them (Fig. 1A). For example, a K6-linked Ub dimer is represented by a tree with two nodes, *V* = {*Ub*_0_, *Ub*_1_}, and one edge, 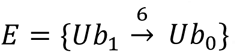. A branched K6/K63-linked trimer is described by *V* = {*Ub*_0_, *Ub*_1_, *Ub*_2_} and 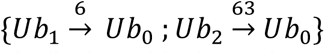 (Fig. 1A). Polyubiquitinated substrates are represented similarly, with the substrate serving as the root node of the tree (Fig. 1A). This representation can accommodate Ub conjugates of arbitrary size, linkage complexity, and topology.

**Figure 1.**
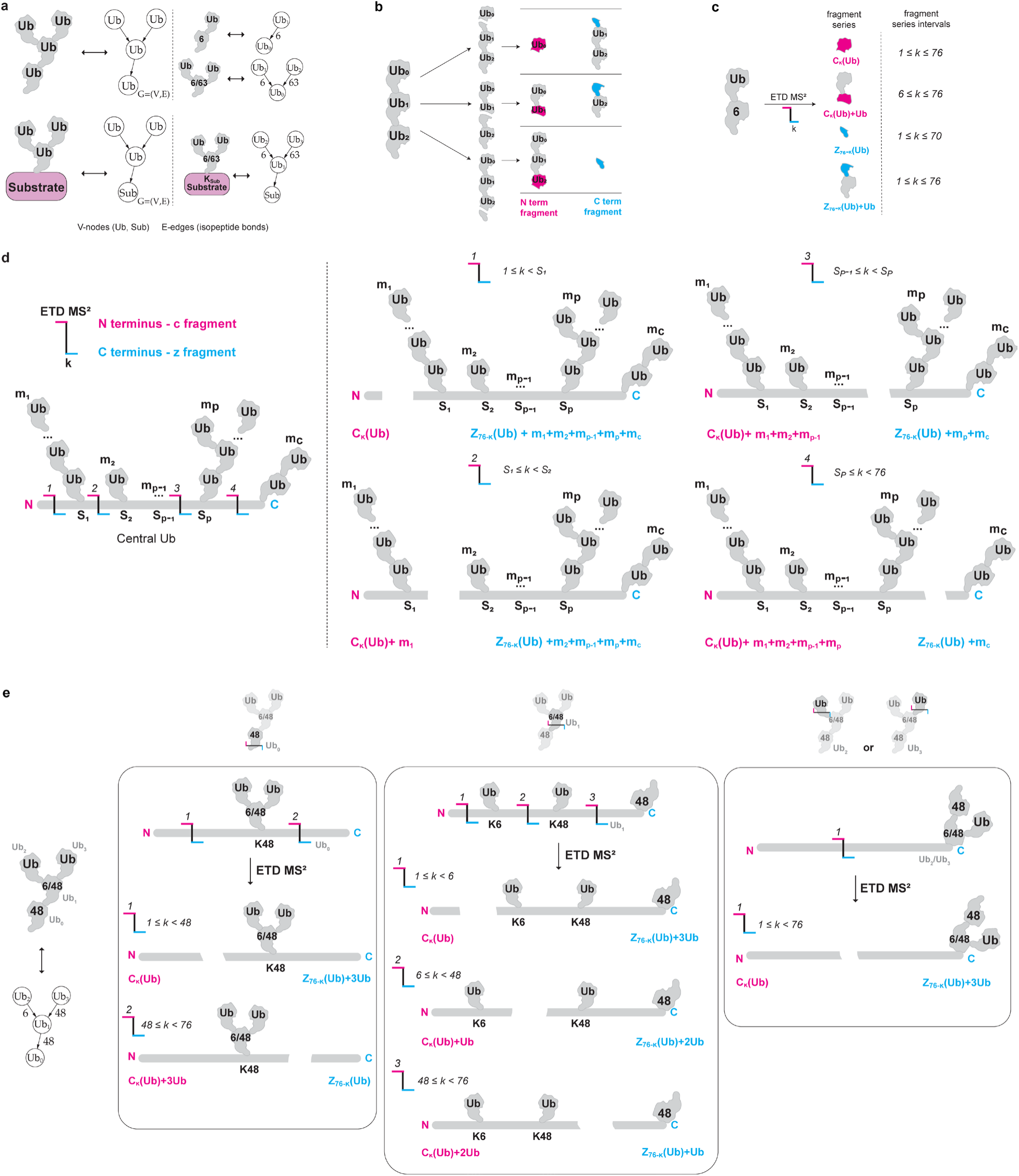
Enumeration of theoretical fragments for Ub conjugates. *(a)* Graph-based representation of Ub conjugates. *(b)* Fragmentation of Ub chain takes place in any of Ub subunits, forming pairs of N-terminal (pink) and C-terminal (cyan) fragments. The fragment series formed by any Ub chain can be described as N- or C-terminal fragment of a single Ub subjected to fragmentation (the part colored in pink or cyan) plus N-number of intact Ub subunits (in grey). *(c)* The example of fragment series and their corresponding fragment series intervals formed by K6 di-Ub upon ETD MS^2^ fragmentation. *(d)* The Ub subunit subjected to fragmentation is named “central”. The whole Ub chain can be represented as a central Ub subunit conjugated to the intact Ub subunits. In the example, central Ub (enlarged in the figure), containing *p* number of Ub sites and *m*_*k*_ intact Ub subunits, is shown. Ub sites split the central Ub into four characteristic regions: downstream *S*_1_, between *S*_1_ and *S*_2_, between *S*_*p*−1_ and *S*_*p*_, and downstream *S*_*p*_ site. Single fragmentation event generates certain fragment series depending on which residue *k* in the central Ub fragmentation occurs. *(e)* Calculation of a total set of fragments for K6/K48/K48 distal tetra-Ub. The chain consists of four (*Ub*_0_, *Ub*_1_, *Ub*_2_, *Ub*_3_) Ub subunits. Fragmentation of *Ub*_0_ occurs downstream or upstream K48-, resulting in two different pairs of fragment series. Fragmentation of *Ub*_1_ can occur downstream of K6-, between K6- and K48-, and downstream K48-, forming three pairs of fragment series. *Ub*_2_ and *Ub*_3_ each generate a single pair of fragment series, as they do not contain conjugated lysine residues. By overlaying the fragment series and their corresponding intervals for all Ub subunits, the final comprehensive set of theoretical fragments is constructed.

Next, we developed an algorithm to translate the graph-based representation of Ub chains into a set of theoretical fragments expected from a TD-MS experiment. This set includes all fragment pairs that can arise from a single backbone cleavage event, depending on the MS fragmentation method used—such as *b/y* ions for collision-induced dissociation (CID) or *c/z* ions for electron transfer dissociation (ETD). Under the single-cleavage assumption, each MS² fragment consists of a partial fragment of one Ub subunit covalently fused to a set of intact Ub subunits (Fig. 1B). For example, ETD of a K6-linked dimer can generate four fragment series: *c*_*k*_(*Ub*) for 1 ≤ *k* < 76; *z*_76−*k*_(*Ub*) for 1 ≤ *k* < 70; *c*_*k*_(*Ub*) + *Ub* for 6 ≤ *k* < 76 and *z*_76−*k*_(*Ub*) for 1 ≤ *k* < 76, where *k* is the residue number at which fragmentation occurs. We define each group of related fragments as a fragment series, and the corresponding *k* values for each series as fragment series intervals (Fig. 1C). Based on the single-cleavage model, the total set of theoretical fragments for a given Ub chain is generated by:(1) simulating cleavage at every residue of each Ub subunit and determining the number of intact Ub subunits that remain fused to the cleaved fragment; and (2) aggregating all resulting fragment series and their intervals across the entire conjugate.

When considering the fragmentation of individual Ub subunits within a chain, the subunit undergoing cleavage (denoted *V*_*i*_) is treated as the central subunit, while the remainder of the chain is represented as a collection of intact Ub subunits covalently attached to it. Figure 1D illustrates a general case in which the central subunit contains *p* ubiquitination sites, each conjugated to *m*_*k*_ intact Ub subunits (where *k* = 1 *to p*), along with a C-terminal carboxyl group linked to *m*_*c*_ additional Ub subunits. The specific values of *m*_*k*_ and *m*_*c*_ depend on the topology of the Ub chain and can be computed using graph search algorithms applied to the oriented tree representation of the Ub chain ^39^.

If we consider that ubiquitination sites divide the central Ub subunit into distinct regions, we can define four fragmentation zones: (1) upstream of site *S*_1_, (2) between *S*_1_ and *S*_2_, (3) between *S*_*p*−1_ and *S*_*p*_, and (4) downstream of *S*_*p*_. Each MS² fragment generated within these regions consists of a fragment of the central Ub subunit (e.g., *c* and *z* ions in ETD) fused to a specific number of intact Ub subunits, depending on chain topology. For example:

1. Upstream of *S*_1_ (1 ≤ *k* < *S*_1_;): *c*_*k*_ and *z*_76−*k*_ + *m*_1_ + *m*_2_ + *m*_*p*−1_ + *m*_*p*_ + *m*_*c*_
2. Between *S*_1_ and *S*_2_ (*S*_1_ ≤ *k* < *S*_2_): *c*_*k*_ + *m*_1_ and *z*_76−*k*_ + *m*_2_ + *m*_*p*−1_ + *m*_*p*_ + *m*_*c*_
3. Between *S*_*p*−1_ and *S*_*p*_ (*S*_*p*−1_ ≤ *k* < *S*_*p*_): *c*_*k*_ + *m*_1_ + *m*_2_ + *m*_*p*−1_ and *z*_76−*k*_ + *m*_*p*_ + *m*_*c*_
4. Downstream *S*_*p*_ (*S*_*p*_ ≤ *k* < 76): *c*_*k*_ + *m*_1_ + *m*_2_ + *m*_*p*−1_ + *m*_*p*_ and *z*_76−*k*_ + *m*_*c*_

Here, *k* denotes the residue at which cleavage occurs, *p* is the total number of modified lysines, and the *m* −values represent the number of intact Ub subunits attached at each modification site or the C-terminus. Given the positions of the ubiquitination sites and the associated *m* −values—each of which can be derived from the Ub chain graph using vertex connectivity analysis—it is possible to enumerate all theoretical fragment series associated with a given central subunit.

To demonstrate this, Figure 1E shows the application of our algorithm to the distal branched K6/K48/K48-linked Ub tetramer. This chain is represented as an oriented tree *G* = (*V*, *E*), where *V* = {*Ub*_0_, *Ub*_1_, *Ub*_2_, *Ub*_3_} and 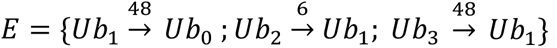. Fragmentation of each Ub subunit is considered separately. For *Ub*_0_, cleavage can occur in two regions:

1. Upstream of K48: *c*_*k*_(*Ub*) and *z*_76−*k*_(*Ub*) + 3*Ub* for 1 ≤ *k* < 48
2. Downstream of K48: *c*_*k*_(*Ub*) + 3*Ub* and *z*_76−*k*_(*Ub*) for 48 ≤ *k* < 76

For *Ub*_1_, cleavage can occur in three regions:

1. Upstream of K6: *c*_*k*_(*Ub*) and *z*_76−*k*_(*Ub*) + 3*Ub* for 1 ≤ *k* < 6
2. Between K6 and K48: *c*_*k*_(*Ub*) + *Ub* and *z*_76−*k*_(*Ub*) + 2*Ub* for 6 ≤ *k* < 48
3. Downstream of K48: *c*_*k*_(*Ub*) + 2*Ub* and *z*_76−*k*_(*Ub*) + *Ub* for 48 ≤ *k* < 76

As *Ub*_2_ or *Ub*_3_ do not contain modified lysines, fragmentation produces only unbranched series: *c*_*k*_(*Ub*) and *z*_76−*k*_(*Ub*) + 3*Ub* for 1 ≤ *k* < 76.

By matching the identical fragment series and unifying fragmentation intervals, the complete set of fragments for K6/K48/K48 can be identified: *c*_*k*_(*Ub*) for *k* ∊ [1,76); *z*_76−*k*_(*Ub*) for *k* ∊ [1,28); *c*_*k*_(*Ub*) + *Ub* for *k* ∊ [6,48); *z*_76−*k*_(*Ub*) + *Ub* for *k* ∊ [1,28); *c*_*k*_(*Ub*) + 2*Ub* for *k* ∊ [48,76); *z*_76−*k*_(*Ub*) + 2*Ub* for *k* ∊ [28,70); *c*_*k*_(*Ub*) + 3*Ub* for *k* ∊ [48,76) and *z*_76−*k*_(*Ub*) + 3*Ub* for *k* ∊ [1,76).

A key strength of our algorithm is that it relies solely on the positions of modified sites and does not require any prior assumptions about the Ub sequence. This generalization allows for accurate prediction of fragmentation patterns even when some vertices in the chain represent non-canonical subunits, such as Ub mutants or ubiquitinated substrates. This flexibility makes our method a universal and robust solution for predicting TD-MS fragmentation patterns across all types of Ub-conjugated proteins.

### Automatic calculation of fragmentation patterns for Ub conjugates

To enable automated interpretation of TD-MS data for Ub conjugates, we developed a Python-based software tool named *UbqTop*. A core component of *UbqTop* is a fragment calculator that uses the algorithm described above to compute theoretical fragment series and corresponding fragment series intervals for any Ub conjugate provided by the user.

To define Ub conjugates in a machine-readable format, we leveraged the mathematical equivalence between oriented trees and linear bracket sequences. In this representation, a Ub conjugate is encoded as a nested bracketed string that captures the topology of the chain. For instance, if Ub subunit *V*_0_ is connected via an isopeptide bond at lysine *S*_1_ to subunit *V*_1_, which is in turn linked at lysine *S*_2_ to subunit *V*_2_, the corresponding bracket sequence is written as: *V*_0_, *S*_1_(*V*_1,_*S*_2_(*V*_2_)). A K6 Ub dimer is encoded as *Ub*, 6(*Ub*), while a branched K6/K63 trimer is represented as *Ub*, 6(*Ub*), 63(*Ub*) (Supplementary Fig.1).

To generate a library of theoretical fragments, the user must:

1. Input the amino acid sequences and assign aliases to each subunit (e.g., wild-type Ub or mutant variants).
2. Provide a list of Ub conjugates in the bracketed format.
3. Select the types of fragments to be annotated (e.g., *c*, *z* ions for ETD).

This information can be entered directly into *UbqTop* via a text interface or uploaded as a formatted text file (Fig. 2A, Supplementary Fig.2, *UbqTop manual*). The software then calculates the complete set of theoretical fragment series and their intervals for each input conjugate. Results are returned in a fragment series table, where columns represent individual Ub conjugates, rows correspond to fragment series, and cells contain the associated fragment series intervals (Fig. 2A, Supplementary Fig.2, *UbqTop manual*). Because *UbqTop* performs analytical calculations without iterating through polypeptide sequences, fragment tables are generated instantaneously, regardless of input library size.

**Figure 2.**
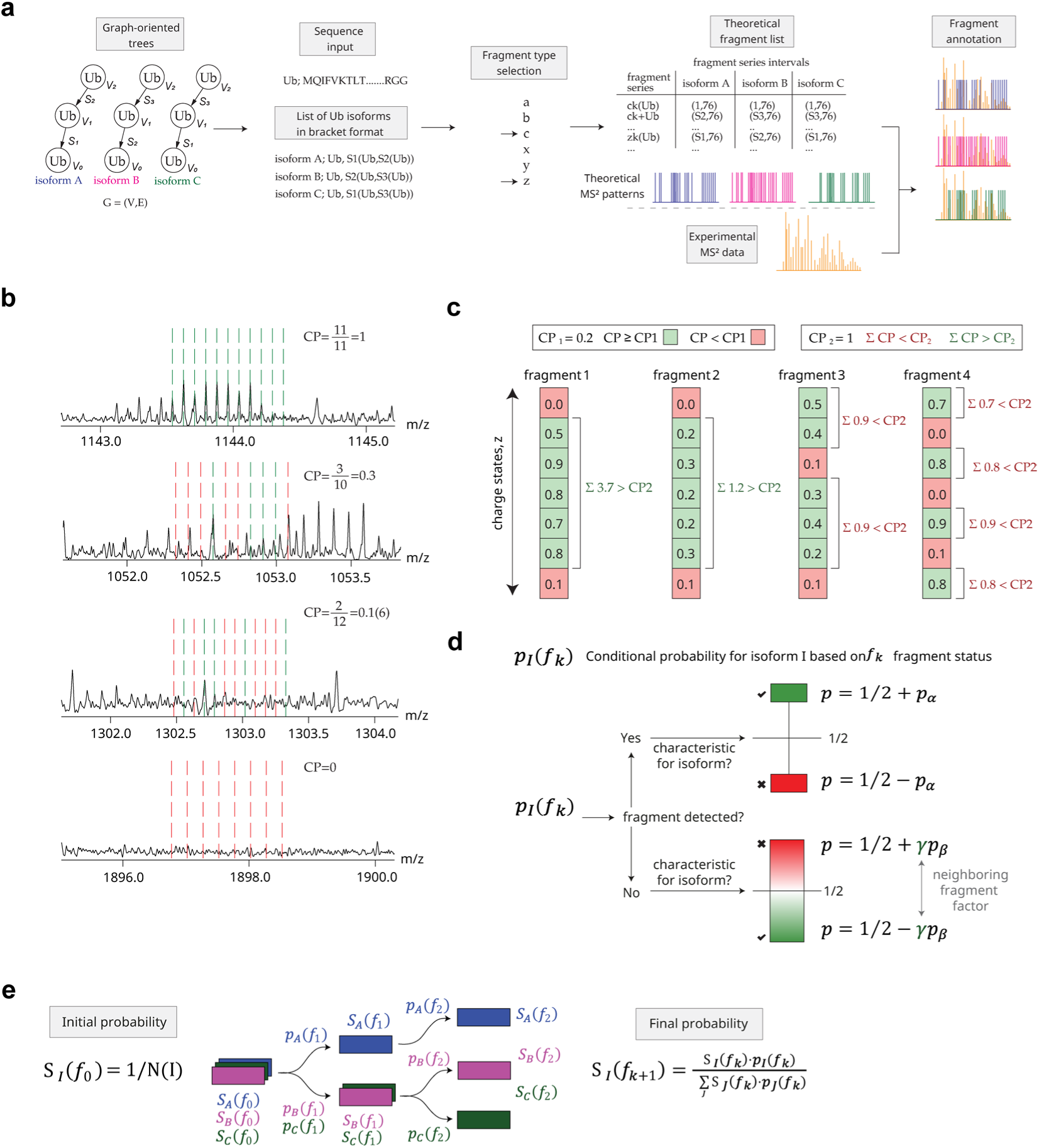
Algorithm for mapping fragmentation data and Bayesian-like approach to calculate probability scores of Ub conjugates. (a) The schematic workflow of data analysis using *UbqTop*. *(b)* Calculation of CP values for four different fragments at single charge states are shown. *(c)* Every fragment at each charge state has the CP value (calculated as described in b), and the values are shown as single cells in a column. Based on the threshold CP_1_ the quality of a fragment at single charge state is assigned (green cell CP>CP_1_, red cell CP<CP_1_). To conclude if the fragment is detected or not, the CP_2_ value is compared with the sum of CP values for consecutively good quality or “green” fragments. If the sum is >CP_2_ then the fragment is detected, if the sum is <CP_2_ the fragment is not detected. *(d)* Calculation of conditional probabilities *p*_*I*_(*f*_*k*_) for the isoform *I* based on the *f*_*k*_ fragment status. *(e)* Calculation of the final probability scores *S*_*I*_(*f*_(*k*+1)_). The initial probability (before fragment detection) *S*_*I*_(*f*_(0)_) is equal for all isoforms in the library and can be calculated as 1/N, where N-is a number of isoforms in the library. The final probability is recalculated for each analyzed fragment with the consideration of conditional probabilities *p*_*I*_(*f*_*k*_) obtained as described in *(d)* for every single fragment.

For polyubiquitinated substrates, users must include the substrate sequence and assign it an alias in the input file. The substrate is designated as the root node of the bracketed tree. *UbqTop* also supports custom sequences, allowing the incorporation of single-point mutations by modifying the wild-type Ub sequence accordingly. Examples of input files for conjugates containing substrates and Ub mutants are provided in the Supplementary Fig.3. Taken together, *UbqTop* enables rapid and flexible generation of theoretical TD-MS fragment sets for any Ub conjugate library—free or substrate-anchored—supporting custom sequence definitions and fragment types. This tool provides a universal solution for predicting complex fragmentation patterns in top-down MS data.

### Annotation of the MS^2^ spectra using targeted fragment search

A critical step in TD-MS–based identification of Ub conjugates is the annotation of experimental MS² spectra through accurate matching to theoretical fragment predictions. To support this capability, we incorporated a targeted fragment search module into the *UbqTop* platform. This module uses a custom algorithm designed to enhance fragment detection sensitivity and maximize sequence coverage by identifying individual isotopic peaks corresponding to theoretical fragments and validating them using a set of continuity-based criteria. Unlike conventional deconvolution-based approaches commonly used for TD-MS data annotation, our strategy focuses on targeted, high-confidence detection, particularly suited to the complex fragmentation patterns of Ub conjugates ^38,40^.

The algorithm begins by calculating the isotopic pattern for each theoretical fragment and scanning the experimental MS² spectrum to identify matching isotopic peaks. For each detected fragment, the algorithm determines the length of the longest consecutive series of observed isotopic peaks, independent of intensity. This value is then normalized to the number of theoretically expected isotopes (those with a relative intensity above a defined threshold), yielding a consecutive percentage (CP) score for each fragment–charge pair (Fig. 2B). A fragment is considered confidently detected if: there is a consecutive series of charge states, where (1) every observed charge state exhibit CP value above a user-defined CP_1_ threshold, and (2) the sum of CP values across all charge states exceeds a global CP_2_ threshold (Fig. 2C). This continuity-based filtering strategy is motivated by two empirical observations: (a) in real MS² spectra, the most intense isotopic peaks of a fragment typically appear as a consecutive series, even at low signal-to-noise ratios; and (b) true fragment signals are often observed as a set of consecutive charge states.

This targeted annotation approach offers several advantages. First, it enables rapid analysis—typically within seconds—of MS² data for any Ub conjugate library, as it avoids computationally intensive isotope envelope fitting. Second, by leveraging the continuity of both isotopic peaks and charge states, the algorithm can detect overlapping fragment signals and confidently identify large, low abundance fragments that may otherwise be missed. In particular, for large fragments, individual charge states may yield modest CP values; however, the presence of multiple, sequential charge states with moderate CPs increases confidence in the fragment assignment. These features make the *UbqTop* annotation strategy both sensitive and robust, particularly for complex or low-signal TD-MS datasets.

### Bayesian-like approach to calculate the probability scores for Ub conjugates

Given the complexity of fragmentation patterns in Ub conjugates, an automated scoring system is essential for accurate identification based on MS² data. In standard TD-MS workflows, proteoforms are typically identified by detecting fragment ions that are unique to a specific sequence. However, this strategy cannot be directly applied to Ub conjugates, as many chain topologies lack uniquely diagnostic fragments. For instance, the full set of fragments produced by a K63-linked Ub dimer is a strict subset of those generated by a K6-linked dimer. As a result, no fragment uniquely associated with the K63 dimer would appear that is absent from the K6 dimer spectrum (Table S1). This overlap complicates isomer discrimination. Accurate identification must consider not only the presence of characteristic fragments but also the absence of others. However, the absence of a fragment cannot be interpreted deterministically, as it may simply reflect incomplete fragmentation or low signal intensity rather than true structural absence. These limitations suggest that TD-MS data for Ub conjugates is best interpreted using a probabilistic rather than a deterministic framework.

To address this, we developed a Bayesian-like algorithm for Ub conjugate identification, which we implemented within the *UbqTop* software. In this approach, the presence or absence of each fragment is treated as evidence contributing to the conditional probability that a particular Ub isomer is present (Fig. 2D). After collecting conditional probabilities across all fragments, a final probability score for each candidate conjugate in the input library is computed based on Bayes’ theorem (Fig. 2E). This scoring framework enables robust identification of Ub conjugates, even in cases where no uniquely diagnostic fragments are observed.

To calculate conditional probabilities, we introduced two influence parameters: *p*_*⍺*_ and *p*_*β*_, corresponding to the presence and absence of a fragment, respectively. When a fragment is observed in the spectrum, the conditional probability assigned to any Ub isoform that can generate this fragment is set to 0.5 + *p*_*⍺*_, while isoforms that cannot generate the fragment are assigned a probability of 0.5 − *p*_*⍺*_. A similar approach is applied to missing fragments using *p*_*β*_, but in this case, the possibility that a fragment was simply not detected—due to low abundance or inefficient fragmentation— must also be taken into account.

To address this, we introduced a probabilistic correction that incorporates local fragment continuity. Specifically, we assume that if a series of consecutive fragments is observed, then the absence of a neighboring fragment likely reflects a structural constraint (i.e., the fragment cannot be produced by that isoform) rather than a detection failure. For example, if the fragments *n Ub* + *c*_*k*_, *n Ub* + *c*_*k*+1_, and *n Ub* + *c*_*k*+2_ are all detected, the absence of *Ub*_*n*_ + *c*_*k*+3_ is more likely to be biologically meaningful than stochastic.

To capture this behavior, we defined a fragment density function, *γ*(*f*_*k*_), 0 ≤ *γ* ≤ 1, which quantifies the likelihood that the absence of fragment *f*_*k*_ is meaningful based on the presence of neighboring fragments in the spectrum. Using this function, the conditional probabilities are adjusted to account for isoforms that cannot generate the fragment (0.5 + *γp*_*β*_) and those that can (0.5 − *γp*_*β*_) (Fig. 2D). Empirically, we found that values of *p*_*⍺*_ = 0.05 and *p*_*β*_ = 0.1 provided optimal performance for TD-MS annotation of Ub conjugates, though these parameters can be tuned depending on the fragmentation method and spectral quality.

The final probability score *S*_*I*_(*f*_*k*+1_) for each Ub isoform *I* is computed iteratively using a Bayesian framework. The algorithm initializes each isoform with an equal prior score *S*_*I*_(*f*_0_) and sequentially incorporates conditional probabilities *p*_*I*_(*f*_*k*_) for each fragment *f*_*k*_ using Bayes’ theorem (Fig. 2E). The iteration continues until all fragments have been processed and the final scores are obtained.

One of the major advantages of *UbqTop* software is its interactivity and tunability for analyzing experimental TD-MS data. *UbqTop* provides visualization tools for both fragment annotation and scoring results, including 2D plots of consecutive percentage (CP) values across fragment series and corresponding sequence coverage maps.

These visual outputs allow users to assess data quality and fine-tune the CP_1_ and CP_2_ thresholds to balance sensitivity and specificity in fragment detection. Moreover, *p*_*⍺*_ and *p*_*β*_ values can be adjusted to optimize isoform discrimination. For example, while lowering CP_1_ and CP_2_ thresholds may increase false-positive fragment identifications, reducing the value of *p*_*β*_ can help compensate by down-weighting absent fragments in the final scoring. This interplay between scoring and detection thresholds highlights the value of integrating theoretical fragment generation, MS^2^ annotation, and Bayesian-like scoring into a unified platform.

### Analysis of free polyubiquitin chains using TD-MS and *UbqTop*

To validate the scoring approach implemented in the *UbqTop* software, we analyzed a series of synthetic Ub isomers using TD-MS with electron-transfer/higher-energy collision dissociation (EThcD) fragmentation. The first test group consisted of six Ub dimers, each linked through a different lysine residue, representing the simplest class of polyubiquitin conjugates. Theoretical fragmentation analysis revealed that two fragment series—*c*_*k*_(*Ub*) + *Ub* and *z*_76−*k*_(*Ub*)—exhibited distinct fragment series intervals across different isomers (Supplementary Table 1). Annotation of the experimental spectra using *UbqTop* detected these differences in sequence coverage, and Bayesian scoring correctly assigned the highest probability to the correct Ub dimer in each case (Fig. 3A, Supplementary Fig.4-Fig.5).

**Figure 3.**
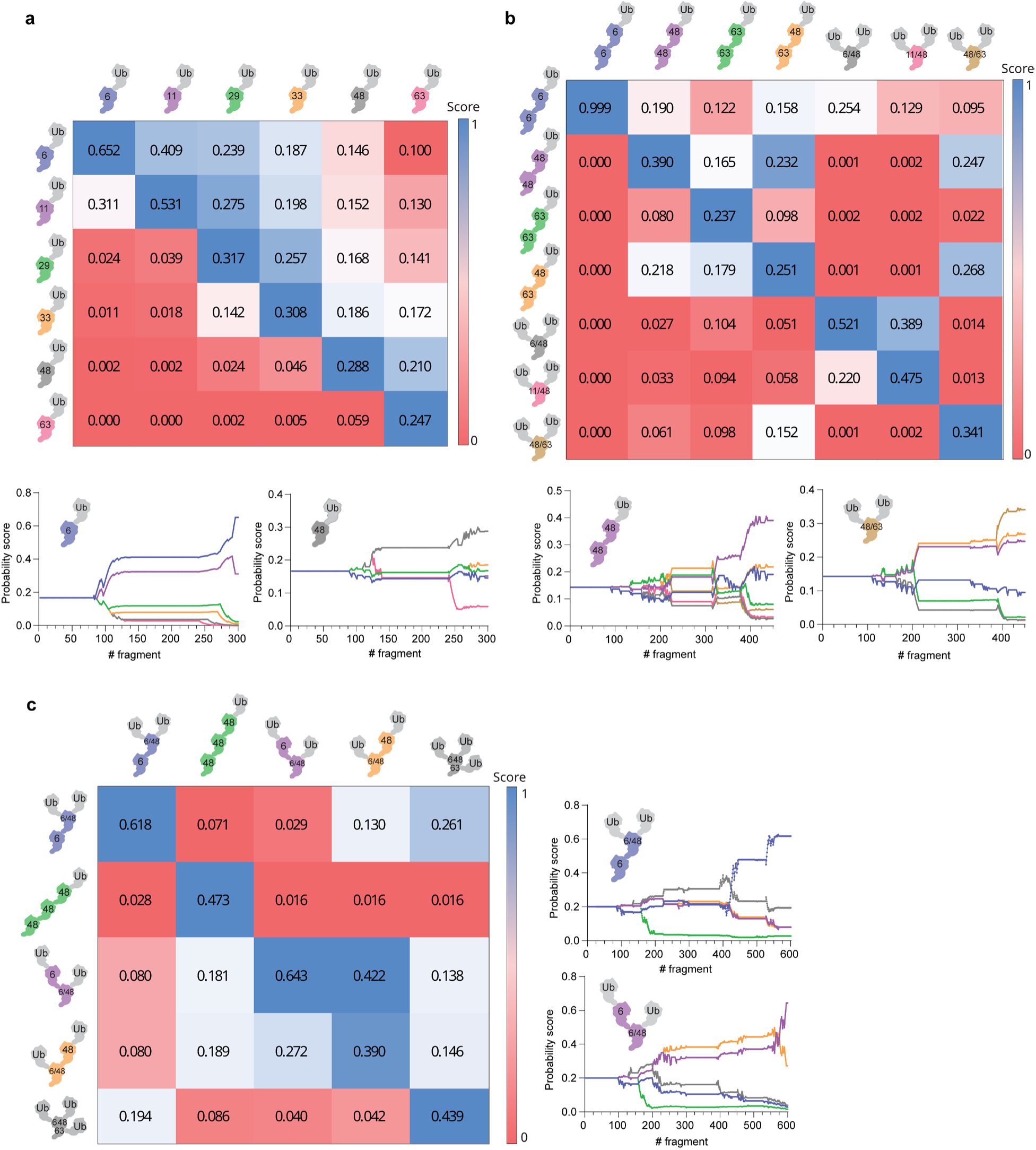
TD-MS analysis and probability scores for free polyubiquitin chains. The scoring results are represented in a table format. Each cell in the table corresponds to a single score of a particular isomer. Dark blue indicates the highest score and the gradient transition to red represents decreasing scores. Examples of the actual scoring plots, exported from *UbqTop*, are shown for several Ub chains. These plots represent the change in probability score (Y axis) in response to each detected or not detected fragment (X axis). (a) Scoring results for six different Ub dimers (K6-, K11, K29-, K33-, K48-, K63-) are shown, with the demonstrated scoring plots for K6-, K48- dimers. (b) Scoring results for seven different Ub trimers (K6- linear, K48-linear, K63-linear, K6/K48 branched, K11/K48- branched, K48/K63- branched, and K63-K48 linear) are shown, with examples of scoring plots for K48- linear, and K48/K63-branched trimers. (c) Scoring results for five different Ub tetramers (K6/K6/K48 distal, K48- linear, K6/K6/K48- proximal, K6/K48/K48- proximal, and K6/K48/K63 triple branched) are shown, with example of scoring plots for K6/K6/K48- distal and K6/K6/K48- proximal tetramers.

In some instances, the differences between scores were less pronounced. For example, for the K63-linked dimer, all isomers received relatively low probability scores (∼0.2), though the correct K63 isomer still achieved the highest score (Fig. 3A, Supplementary Fig. 5.). This can be attributed to the K63 dimer producing the fewest characteristic fragments across its unique series, limiting the available discriminatory signal (Supplementary Table 1). A similar challenge was observed for the K29- and K33-linked dimers, which share highly similar fragment series intervals, resulting in close but distinguishable probability values (Supplementary Table 1, Supplementary Fig. 5.).

Despite these challenges, the Bayesian framework was able to interpret both the presence and absence of characteristic fragments to resolve isomer identity, consistently assigning the highest score to the correct Ub conjugate. These results confirm that *UbqTop* can reliably distinguish Ub isomers using TD-MS data, even in cases where fragments overlap or sparse coverage limits traditional deterministic identification.

Next, we increased the complexity of the Ub conjugates by synthesizing a library of seven Ub trimers with distinct linkage compositions and chain architectures. Compared to dimers, the number of informative fragment series increases for trimers, allowing combinations such as *c*_*k*_(*Ub*) + *Ub*, *c*_*k*_(*Ub*) + 2*Ub*, *z*_76−*k*_(*Ub*), *z*_76−*k*_(*Ub*) + *Ub* —each observed over specific fragment series intervals—to be used for isomer discrimination (Supplementary Table 2). Some series, particularly those involving larger fragments (e.g., *c*_*k*_(*Ub*) + 2*Ub*), may be difficult to detect due to their higher mass and correspondingly low signal-to-noise (S/N) ratio (Supplementary Fig. 6). To account for the presence of the lysine-to-arginine replacements in several Ub chains without compromising the scoring process evaluation, instead of uploading mutant-containing Ub sequences, the sequences of the intact or fragmented portion in the fragment series were modified.

Across all seven Ub trimers, the scoring algorithm correctly assigned the highest probability to the true isomer (Fig. 3B, Supplementary Fig. 7-Fig. 8). As with dimers, K63-linked trimers exhibited lower absolute scores due to the limited number of characteristic fragments, which can reduce scoring resolution (Supplementary Table 2). Notably, the absence of fragments in specific series—such as *c*_*k*_(*Ub*) + 2*Ub* —did not adversely affect scoring outcomes. This demonstrates that the algorithm effectively handles sparse data by down-weighting fragment series with insufficient signal, eliminating the need for manual filtering and reducing the risk of false-positive identifications.

Lastly, we synthesized a library of five Ub tetramers that varied in linkage composition, number of branch points, and the position of the branch point relative to the proximal Ub subunit. These topological differences—particularly the location of branching—are not preserved in middle-down MS workflows, which fragment the chain and lose critical structural context. In contrast, our approach retains this information and enables isomer discrimination at full-chain resolution.

*UbqTop* successfully identified the correct isomer in all five cases (Fig. 3C). In rare instances, however, highly similar isomers yielded nearly identical scores. For example, the K6/K48/K48-proximal tetramer scored almost equivalently to the K6/K6/K48-proximal isomer (Fig. 3C, Supplementary Fig. 9-Fig. 10). This ambiguity arises because these two topologies differ only in the *c*_*k*_(*Ub*) + 2*Ub* and *z*_76−*k*_(*Ub*) + *Ub* fragment series—series that are absent in the K6/K48/K48-proximal isomer, which cannot produce these fragments (Supplementary Table 3). In such cases, accurate discrimination depends entirely on the absence of specific fragments, which reduces scoring resolution.

Nonetheless, the algorithm consistently distinguished topologies such as proximal vs. distal branching, linear vs. branched, and double vs. triple-branched chains. To our knowledge, this is the only method capable of distinguishing Ub chain isomers that differ solely in the position of the branch point—an analytical capability not achievable with middle-down MS.

### Strategy for the analysis of polyubiquitinated substrates using TD-MS

While top-down MS (TD-MS) offers powerful capabilities for analyzing free Ub chains, it faces important limitations when applied to polyubiquitinated substrates—particularly in cases where multiple isomeric Ub chains are conjugated to different sites on the same protein. Under single-fragmentation conditions, cleaving the substrate reveals nothing about the topology of the attached Ub chains, whereas cleaving within the Ub chains obscures information about their site of attachment to the substrate (Supplementary Fig. 11). This disconnect between chain topology and localization presents a fundamental challenge for comprehensive analysis. In principle, such ambiguity could be resolved using MS³ workflows; however, the inherently low yield of MS³ experiments, combined with the lack of standardized or automated data analysis pipelines, severely limits their practical utility for routine characterization.

To determine both the topology of a Ub chain and its precise attachment site on a substrate, we developed a strategy involving selective proteolysis of the substrate while keeping the Ub chain intact. Digestion of the substrate under native conditions reduces the number of lysine residues per peptide of the substrate, improving the resolution of site localization. We screened several proteases and identified Asp-N as particularly well-suited: it efficiently digests protein substrates while exhibiting minimal activity toward Ub chains, enabling simultaneous analysis of chain architecture and site of modification. This stands in contrast to trypsin and LbPro*, which cleave Ub at R74 under native conditions, disrupting chain continuity and chain-site connectivity (Fig. 4A) ^22^. This approach also simplifies TD-MS analysis by substantially reducing the size of the substrate. The resulting Ub–substrate fragments are more amenable to ionization and exhibit efficiencies comparable to free Ub chains, making them well-suited for TD-MS workflows even at chromatographic scale. We validated the resistance of Ub chains to Asp-N cleavage using both low- and high–molecular weight conjugates, across a range of substrate-to-protease ratios and incubation times. We observed minimal cleavage at high protease:substrate ratio upon overnight incubation for high molecular weight Ub chains, with the low molecular weight chain remaining intact under the same conditions (Fig. 4A, Supplementary Fig. 12).

**Figure 4.**
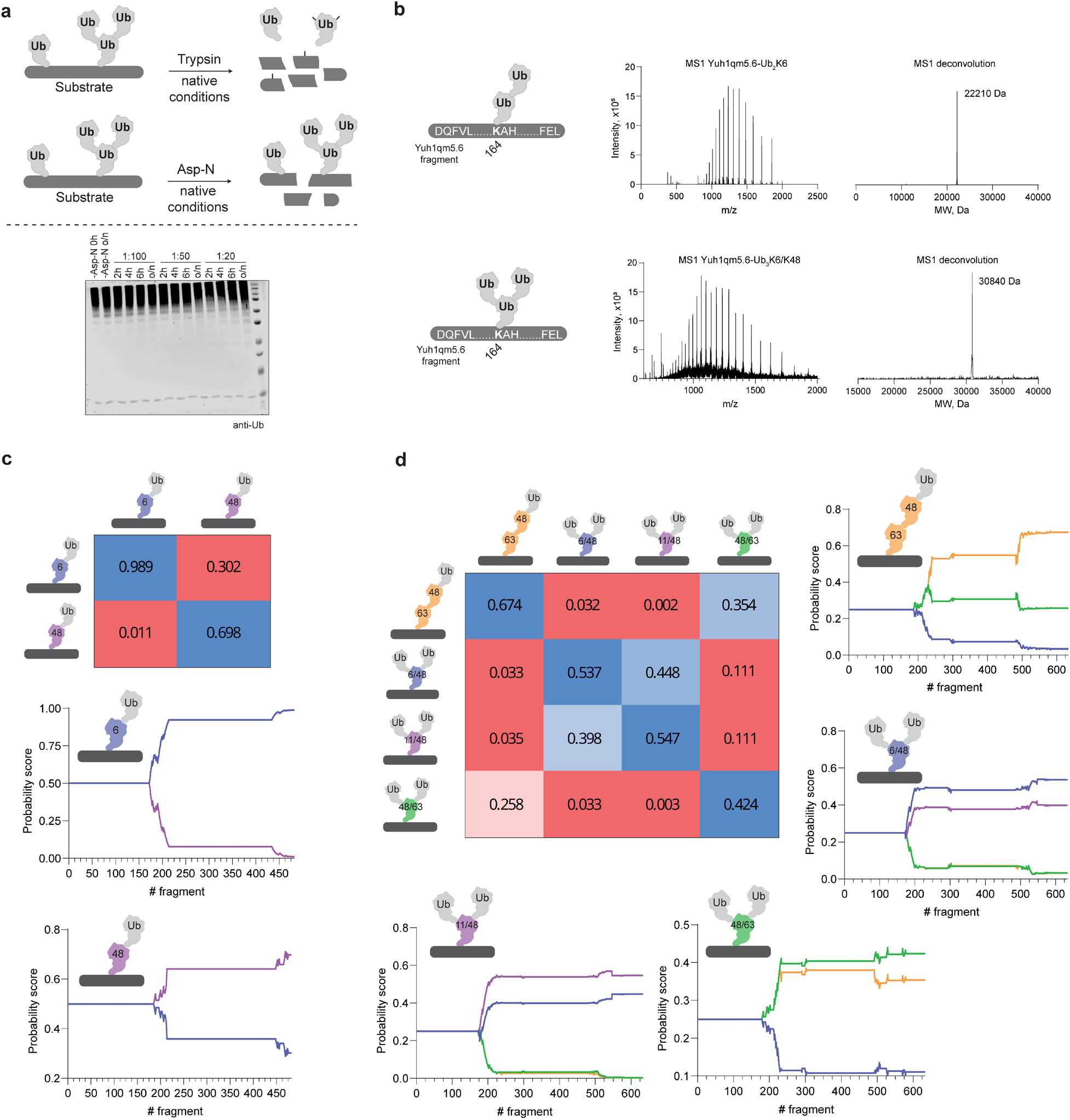
Approach for the analysis of ubiquitinated substrates using TD-MS. (a) Schematic representation of trypsin and Asp-N digestion of ubiquitinated substrates under native conditions. In comparison with trypsin, which cleaves both the Ub chain and a substrate, Asp-N digestion results in the cleavage of a substrate, preserving Ub chain(s) conjugated to the substrate. Anti-Ub blot shows the result of Asp-N digestion of high molecular weight Ub chains under native conditions with different substrate:protease mass ratios and incubation times. (b) Yuh1qm5.6, which has a single Ub site at the K164 residue, was used as a model substrate to test Asp-N activity toward ubiquitinated substrates. Yuh1qm5.6 conjugated to K6-Ub dimer or K6/K48 branched Ub trimer was digested with Asp-N, and MS^1^ analysis was performed to check if the Yuh1qm5.6 fragment conjugated to the intact chain is detected. MS^1^ spectrum deconvolution resulted in 22.2 and 30.84 kDa fragment masses corresponding to the Yuh1qm5.6 fragment fused to the intact Ub dimer and trimer, respectively. (c) Scoring results for di-ubiquitinated Yuh1qm5.6 variants are shown (Yuh1qm5.6 fused to: K6, K48 dimers). (d) Scoring results for tri-ubiquitinated Yuh1qm5.6 variants are shown (Yuh1qm5.6 fused to: K63-K48 linear, K6/K48 branched, K11/K48-branched, and K48/K63-branched trimers).

To evaluate this approach in a controlled setting, we used a model substrate, Yuh1qm5.6—a variant of the C-terminal hydrolase Yuh1 that exhibits enhanced transamidation activity over hydrolysis ^41^. This property enables autoubiquitination of Yuh1qm5.6 at a single, defined lysine residue (K164) with polyubiquitin chains of varying length and topology. We first generated Yuh1qm5.6 conjugated to di- and tri-Ub chains and assessed whether, following Asp-N digestion, the expected substrate fragment remained covalently attached to an intact Ub chain at the MS¹ level. Based on the known Yuh1qm5.6 sequence, the Asp-N–derived substrate fragment was predicted to have a molecular weight of 5.1 kDa. Accordingly, the expected molecular weights for Yuh1qm5.6 conjugated to two and three Ub units were 22.2 kDa and 30.8 kDa, respectively—both of which were clearly detected in the MS¹ spectra (Fig. 4B).

We next asked whether MS² fragmentation under LC-MS conditions would yield sufficient sequence coverage to confidently identify the Ub chain isomer attached to Yuh1qm5.6. Unlike direct infusion, which allows extended ion accumulation times and is commonly used for free chain analysis, LC-MS is required here to remove Asp-N and reaction byproducts but inherently limits accumulation time. Even with this constraint, the MS² data generated from online LC-MS provided sufficient sequence coverage, as illustrated by coverage maps for several characteristic fragment series (Supplementary Table 4-5, Supplementary Fig. 13-Fig. 14).

Finally, we applied the *UbqTop* scoring algorithm to the acquired MS² spectra. As shown in Figure 4C-D, the correct Ub isomer was identified in all Yuh1qm5.6 conjugates, demonstrating that chain topology can be accurately determined from substrate-conjugated Ub species even under the limited acquisition time available in LC-MS workflows. To date, TD-MS analysis has been applied primarily to identify ubiquitination sites on substrates by analyzing monoubiquitinated species, but not to reconstruct the topology of attached polyubiquitin chains ^35^.

### Characterizing site-specific Ub chain architecture on EGFP

To demonstrate the utility of our approach for deciphering complex and previously uncharacterized ubiquitination patterns, we applied it to EGFP, a model substrate with multiple lysine residues. EGFP was ubiquitinated in vitro using the UbcH7/NleL enzyme pair, which primarily generates K6- and K48-linked polyubiquitin chains, with some activity toward K11 and K63 linkages^25,42^. Given the multiple potential ubiquitination sites and chain types, the resulting modification landscape is highly heterogeneous and not amenable to conventional analysis.

We first evaluated the efficiency of Asp-N digestion on non-ubiquitinated EGFP and confirmed complete coverage at the MS¹ level, with all theoretical fragments observed (data is not shown). We then isolated polyubiquitinated EGFP species from the reaction mixture and performed Asp-N digestion under native conditions (Fig. 5A). MS¹ analysis of the digest revealed three distinct EGFP fragments modified with up to two intact Ub subunits (Fig. 5B). Species containing more than two Ub units were likely present but fell below detection thresholds due to low abundance and sample complexity under LC-MS conditions.

**Figure 5.**
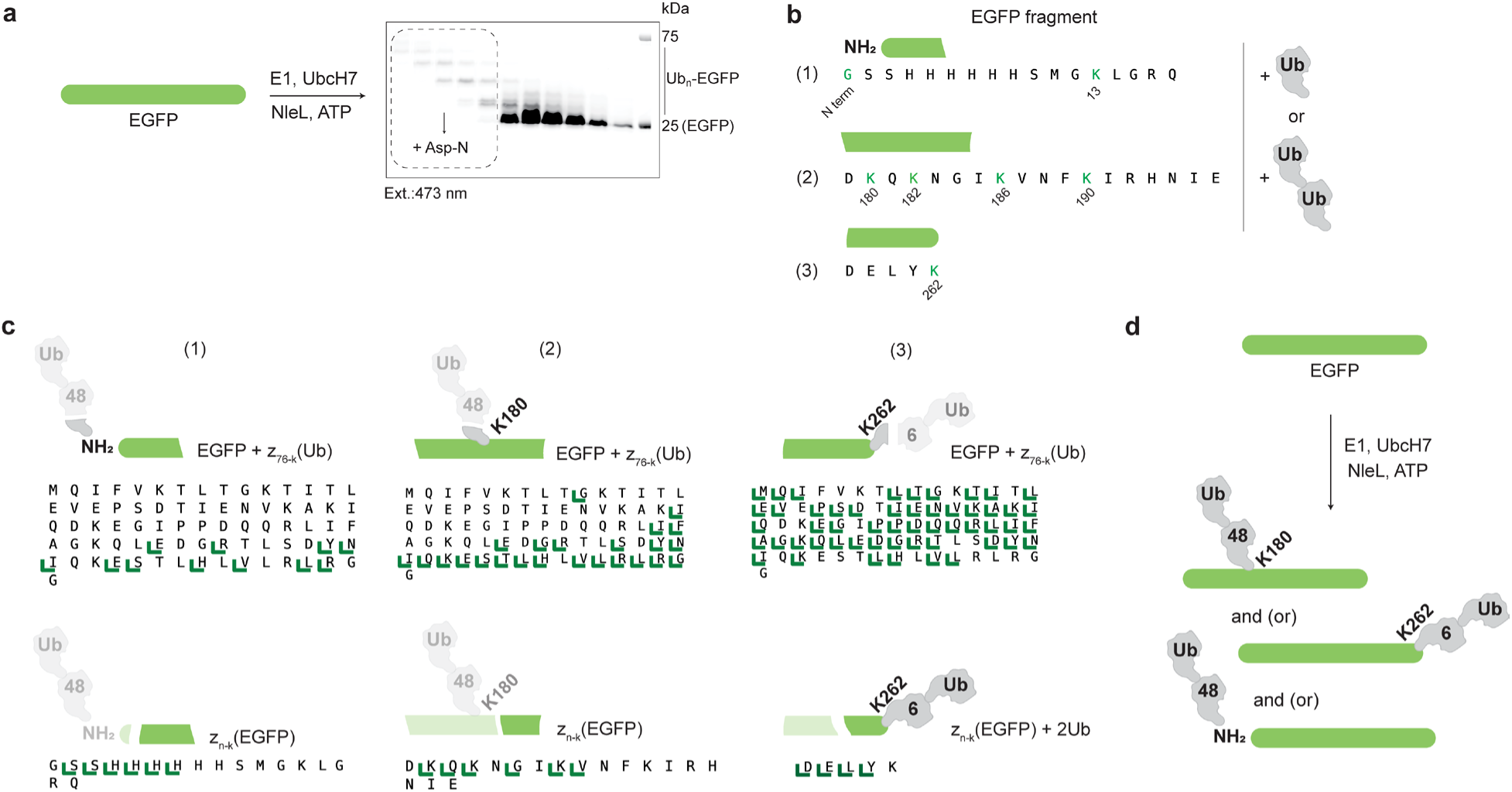
Assessing Ub pattern on EGFP upon UbcH7/NleL enzymatic reaction. (a) Schematic representation of EGFP ubiquitination using E1, UbcH7, and NleL. The fluorescent gel (excitation at 473 nm) from size exclusion purification of Ub_n_-EGFP chains is shown. The dashed rectangle highlights Ub_n_-EGFP chains collected for Asp-N digestion and TD-MS analysis. (b) EGFP fragments modified with one or two intact Ub subunits identified on MS^1^ level after Asp-N digestion (1-3). The highlighted residues indicate potential sites for ubiquitination. (c) TD-MS analysis of EGFP fragments modified with two intact Ub subunits. EGFP+z_76-k_(Ub) fragments series coverage maps are shown, that were primarily used to identify Ub chain type anchored on each fragment. z_n-k_(EGFP) or z_n-k_(EGFP)+2Ub were used to identify the Ub site on EGFP. (d) Schematic representation of the whole Ub pattern of EGFP recapitulated from TD-MS data.

Each fragment contained one or more potential ubiquitination sites: fragment 1 could be modified at K13 or the N-terminus; fragment 2 contained four lysines (K180, K182, K186, K190); and fragment 3 included a single lysine (K262). In the case of di-ubiquitination, modifications could reflect either a di-Ub chain at a single site or two monoubiquitination events.

To resolve both Ub chain type and site of attachment, we performed ETD-MS² analysis on all three fragments and analyzed the data using *UbqTop*. We prioritized fragment series with sufficient detection and isoform-specific fragment intervals (Supplementary Table 6). Fragment 1 was found to be modified at the N-terminal glycine—not K13— based on the observed *z*_*n*−*k*_(*EGFP*) fragment coverage. The attached chain was identified as a K48-linked di-Ub, supported by *EGFP* + *z*_76−*k*_(*Ub*) coverage spanning residues 1–28, consistent with the expected fragment series for a K48 dimer (Fig 5C, Supplementary Table 6, Supplementary Fig. 15).

For fragment 2, analysis of *z*_*n*−*k*_(*EGFP*) ions pinpointed K180 as the predominant site of modification, and the conjugated chain was again identified as a K48 dimer. Fragment 3, containing a single lysine (K262), was also unambiguously assigned a K6-linked di-Ub, confirmed by a *EGFP* + *z*_76−*k*_(*Ub*) fragment coverage interval of 6–76, matching the theoretical pattern for K6 linkages (Fig 5C, Supplementary Table 6, Supplementary Fig. 15). Notably, the site-specific identification of chain type and conjugation site was possible without any prior knowledge of Ub site occupancy or linkage composition— information that cannot be resolved by traditional proteolysis-based methods.

To benchmark our findings, we analyzed the same ubiquitinated EGFP pool using bottom-up proteomics. Trypsin digestion under denaturing conditions confirmed the presence of K6- and K48-linked Ub chains, in agreement with our TD-MS results (Supplementary Table 7, Supplementary Fig. 16). We also detected GG-modified peptides corresponding to all three Ub sites identified by TD-MS (N-terminus, K180, K262), as well as additional low-abundance modifications at K182, K186, and K190. One additional site, K125, was detected only by bottom-up MS (Supplementary Table 8). Its relatively high abundance suggests that it may be modified by longer or more heterogeneous chains that were below the detection limit of the TD-MS workflow.

Together, these results demonstrate the power of our top-down strategy combined with *UbqTop* analysis to characterize, for the first time, the precise architecture of Ub chains at specific sites on a substrate protein. This integrated approach uniquely resolves both chain type and site of attachment in a single experiment—information that is inherently lost in bottom-up or middle-down workflows. As shown with EGFP, this capability enables high-resolution decoding of complex ubiquitination patterns, including those with mixed linkage types and multiple modification sites.

## DISCUSSION

Here, we present an integrated top-down mass spectrometry (TD-MS) approach for decoding the architecture and site-specific positioning of Ub chains on substrates. Central to this strategy is *UbqTop*, a software platform that automatically generates theoretical fragment libraries, performs targeted annotation of MS² spectra, and calculates probabilistic scores for Ub conjugate identification. *UbqTop* was validated across a range of Ub chain types—free and substrate-anchored—including isomeric chains that differ only in branching patterns or linkage topology. The method achieves high sensitivity in distinguishing these species, even when fragment differences are subtle or defined solely by the absence of specific ions.

To enable precise mapping of Ub chains to individual sites on protein substrates, we incorporated a native proteolysis step using Asp-N, which cleaves the substrate while preserving Ub chains. This approach circumvents the need for MS³ and allows site-specific chain identification directly at the MS² level. Starting from a model substrate with a single known Ub site (Yuh1qm5.6) and progressing to a complex, multiply ubiquitinated substrate (EGFP) with unknown modification patterns, we show that *UbqTop* can resolve both chain type and conjugation site—a level of detail not attainable with bottom-up or middle-down proteomics.

A key advance of *UbqTop* is its automation of theoretical fragment generation and scoring, eliminating the need for manual curation of fragmentation patterns. The annotation algorithm incorporates charge-state and isotopic continuity filters to reliably detect low-intensity fragments without inflating false positives. Parameter settings— such as fragment quality thresholds and influence weights for detected or missing ions—can be tuned to accommodate varying data quality, making the software robust across instruments and acquisition platforms.

The scoring algorithm uses a Bayesian-like framework that integrates both the presence and absence of ions to compute probabilities for each Ub isomer. This feature is especially powerful for isomeric chains that differ only in fragment series intervals, where the absence of specific fragments is critical to correct identification.

A key limitation of TD-MS—shared across most current implementations—is its inability to deconvolute mixtures of isomeric Ub chains. When multiple isomers co-elute, their fragmentation spectra merge, complicating unambiguous identification. Conventional reverse-phase chromatography generally lacks the resolving power to separate such species, but emerging techniques like ion mobility spectrometry (IM-MS) may offer a viable solution. Additionally, when a single Ub site on a substrate is modified by multiple chain types, the resulting spectral overlap precludes confident assignment of isomers. Overcoming these challenges will require further advances in front-end separation and acquisition strategies, such as fragmentation at MS^3^ level.

We also acknowledge the general risk of false-positive identifications inherent to any computational scoring method. While default *UbqTop* parameters performed consistently across tested datasets, the software offers extensive user control over all critical inputs—from *m/z* tolerance to scoring thresholds—and provides tabular and graphical outputs at every stage to support transparency and manual validation.

In sum, this work demonstrates the first automated, end-to-end TD-MS platform capable of characterizing Ub chain architecture and attachment sites across diverse substrates. *UbqTop* represents a significant advance over existing proteomics workflows by preserving full information about Ub chain topology and site specificity. We anticipate that this approach will expand access to TD-MS for Ub research and accelerate mechanistic studies of Ub signaling. Future advances in MS hardware and separation methods will further enhance the resolution and scope of this platform, enabling the routine analysis of complex, high–molecular-weight ubiquitinated species.

## MATERIALS AND METHODS

### Protein expression and purification

*Wild-type Ub along with lysine-to-arginine variants* were expressed in E. coli Rosetta 2(DE3)pLysS cells and purified by perchloric acid precipitation, following procedure adapted from ref.^43^

*E1, UBE2R1 (cdc34), UBE2N/UBE2V2 (UbcH13-MmS2)* were expressed in Rosetta 2(DE3)pLysS E.coli cells in LB media supplemented with appropriate antibiotics at 37°C to OD600 0.6-0.8 and after induction with IPTG incubated at 18°C for 16 h. Cultures were harvested at 5,000g for 20 min 4°C, resuspended in lysis buffer A (50 mM Tris pH 7.5, 300 mM NaCl, 1 mM TCEP and 10 mM imidazole), lysed by sonication, and clarified by centrifugation at 35,000g for 30 min 4°C. Clarified lysate was then incubated with Ni-NTA resin for 2 h, washed with lysis buffer A, and eluted into Ni-NTA elution buffer (lysis buffer A plus 300 mM imidazole).

*Ube2L3 (UbcH7) and UBE2S-UBD* constructs were expressed in Rosetta2(DE3)pLysS E.coli cells in LB media supplemented with appropriate antibiotics at 37°C to OD600 0.6-0.8 and transferred to 16°C for 16 h after induction with IPTG. Cultures were harvested at 5,000g for 20 min 4°C, resuspended in lysis buffer B (270 mM sucrose, 50 mM Tris pH 8.0, 50 mM NaF, and 1 mM DTT), lysed by sonication, and clarified by centrifugation. Clarified lysate was then incubated with GST resin for 2 h, washed with high salt buffer (25 mM Tris pH 8.0, 500 mM NaCl, and 5 mM DTT) and then by low salt buffer (25 mM Tris pH 8.0, 150 mM NaCl, and 5 mM DTT), and resuspended in 3C protease buffer (50 mM Tris pH 8.0 and 150 mM NaCl) for on-resin cleavage with HRV 3C protease overnight or thrombin cleavage buffer (30 mM Tris pH 7.5, 150 mM NaCl, 2.5 mM CaCl2) for on resin thrombin cleavage.

*NleL (aa 170-782) or NleL E705A* was expressed in BL21(DE3)pLysS E.coli cells in LB media supplemented with appropriate antibiotics at 37°C to OD600 0.6-0.8 and transferred to 16°C for 16 h after induction with IPTG. Cultures were harvested at 5,000g for 20 min 4°C, resuspended in lysis buffer C (50 mM Tris pH 8.0, 200 mM NaCl, 1 mM EDTA and 1 mM DTT), lysed by sonication, and clarified by centrifugation. Clarified lysate was then incubated with GST resin for 2 h, washed with lysis buffer, and eluted into GST elution buffer (lysis buffer C plus 10 mM reduced glutathione). The eluate was concentrated and buffer exchanged into TEV protease buffer (50 mM Tris pH 8.0, 150 mM NaCl, and 0.5 mM TCEP) and cleaved overnight with TEV protease. The 6xHis-tagged Yuh1 and His-EGFP were expressed in BL21(DE3) E. coli. The cells were lysed in 50 mM Tris pH 8.0, 300 mM NaCl, 10 mM imidazole, and purified using nickel (Clontech) affinity chromatography. The equilibration buffer was 50 mM Tris pH 8, 300 mM NaCl, 10 mM imidazole and the elution buffer was 50 mM Tris pH 8.0, 300 mM NaCl, 300 mM imidazole.

All the proteins were further purified using size exclusion chromatography on Superdex75 or Superdex200 (GE) with the running buffer 50 mM Tris pH 7.5 (or pH 8.0), 150 mM NaCl, and 1 mM DTT, 5% glycerol. Some proteins were purified using anion exchange chromatography using 20 mM Tris pH 8.0, 1 mM DTT as Buffer A and 20 mM Tris pH 8.0, 1M NaCl, 1 mM DTT as Buffer B.

### Generation of native Ub chains

#### Ub dimers

*K48-linked ubiquitin dimer:* 1 mM Ub, 0.3 uM E1, and 5 uM Cdc34 were mixed in reaction buffer A (20 mM ATP, 10 mM MgCl2, 40 mM Tris-HCl pH 7.5, 50 mM NaCl, and 1.5 mM DTT) and incubated for 6 hours at 37°C. *K63-linked ubiquitin dimer:* 0.5 mM Ub, 1.5 uM E1, and 0.75 uM Ubc13-Mms2 were mixed in reaction buffer A and incubated at 37 for 9 h. K6-linked ubiquitin dimer: 2 mM Ub, 1.5 uM E1, 10 uM UbcH7, and 1uM NleL were mixed in reaction buffer A. 5 uM OTUB1 and 3 uM AMSH were added after 3 h of incubation and the mixture was left overnight at 37°C. All reactions were quenched with 50 mM ammonium acetate pH 4.4. All synthesized ubiquitin dimers were further purified from the reaction mixture using size exclusion chromatography (Superdex75) using buffer B (50 mM Tris-HCl pH 7.5, 300 mM NaCl). The generated ubiquitin dimers were buffer exchanged into MiliQ water and stored in a lyophilized form until needed. *K11-, K29- and K33-linked ubiquitin* dimers were purchased from R&D systems Inc. (Minneapolis, MN). Concentrations of all ubiquitin dimers were measured by BCA assay (Thermo Scientific).

#### Ub trimers

*K48-linked tri-Ub:* 2 mM Ub, 1 uM E1, and 10 uM cdc34 were mixed in reaction buffer A (20 mM ATP, 10 mM MgCl_2_, 40 mM Tris-HCl pH 7.5, 50 mM NaCl, and 1.5 mM DTT) and incubated for 6 h at 37°C. *K63-linked tri-Ub:* 1 mM Ub, 1.5 uM E1, and 0.75 uM Ubc13-Mms2 were mixed in reaction buffer A and incubated at 37°C for 9 h. *K6-linked tri-Ub:* 2 mM Ub, 0.5 uM E1, 10 uM UbcH7, and 1uM NleL were mixed in reaction buffer A. 5 uM OTUB1 and 3 uM AMSH were added after 3 h of incubation and the mixture was left overnight at 37°C. *K6/K48 branched triUb:* 2 mM Ub K6R/K48R, 1 mM Ub D77, 0.5 uM E1, 10 uM UbcH7, 1 uM NleL, 3 uM AMSH were mixed in reaction buffer A and incubated overnight at 37 °C. *K48/K63-branched tri-Ub:* 2 mM Ub K48R/K63R, 1 mM Ub D77, 0.5 uM E1, 10 uM cdc34, 1uM UbcH13-MmS2 were mixed in reaction buffer A and incubated overnight at 37 °C. *K11/K48-branched tri-Ub:* 2 mM Ub K11R/K48R, 1 mM Ub D77, 0.5 uM E1, 10 uM cdc34, 20 uM UBE2S-UBD were mixed in reaction buffer A and incubated overnight at 37 °C. *K63-K48-linear tri-Ub:*1) 1.5 mM Ub K48R D77, 1.5 mM Ub K63R, 0.5 uM E1, 0.75 uM UbcH13-MmS2 were mixed in reaction buffer A and incubated at 37°C overnight. The resulting Ub dimer was purified using size exclusion chromatography (Superdex75) and buffer exchanged into MiliQ water. 2) 0.5 mM of Ub dimer purified in step (1) was mixed with 0.5 mM Ub K48R, 3 uM E1, 20 uM cdc34 in reaction buffer A and incubated at 37°C overnight.

For all branched and heterotypic trimers, the enzymatic reaction mixture was further treated with 0.5 uM Yuh1 at room temperature overnight to remove the C-terminal D77 residue from the proximal Ub.

All reactions were quenched with 50 mM ammonium acetate pH 4.4. All synthesized ubiquitin trimers were further purified from the reaction mixture using size exclusion chromatography (Superdex75) using buffer B (50 mM Tris-HCl pH 7.5, 300 mM NaCl) and buffered exchanged into MiliQ water. The concentration was measured using BCA assay (Thermo Scientific).

#### Ub tetramers

*K48-linear tetra-Ub:* 2 mM Ub, 1 uM E1, and 10 uM cdc34 were mixed in reaction buffer A (20 mM ATP, 10 mM MgCl_2_, 40 mM Tris-HCl pH 7.5, 50 mM NaCl, and 1.5 mM DTT) and incubated overnight at 37°C. *K6/K6/K48 proximal tetra-Ub were synthesized in three steps:* 1) 1.5 mM Ub K6R/K48R, 1 mM Ub D77, 0.5 uM E1, 10 uM cdc34, were mixed in reaction buffer A and incubated for 6 hours at 37°C. The dimer fraction was purified on size exclusion chromatography (Superdex 75) using buffer B (50 mM Tris-HCl pH 7.5, 300 mM NaCl), and buffer exchanged into MiliQ water. 2) 2 mM Ub K6R/K48R, 1mM Ub K48R D77, 10 uM UbcH7, 1 uM NleL E705A were mixed in reaction buffer A and incubated overnight at 37°C. The dimer fraction was purified as described in step 1 and treated with Yuh1 to remove the proximal D77 residue. 3) The dimers from step 1) and 2) were mixed in the equimolar ratio (50 uM) with 3 uM E1, 10 uM UbcH7, 1uM NleL, 3uM AMSH in reaction buffer A and incubated overnight at 37°C. *Ub K6/K48/K48 proximal were generated in three steps:* 1) K6/K48 branched trimer (synthesized from Ub K6R/K48R and Ub D77) with the intact D77 on a proximal subunit was treated with 3 uM OTUB1 for 6 hours at 37°C. The resulting dimer was purified using size exclusion chromatography using buffer B and buffer exchanged into MiliQ water. 2) 1.5 mM Ub K6R/K48R, 1 mM Ub D77, 0.5 uM E1, 10 uM cdc34, were mixed in reaction buffer A and incubated for 6 hours at 37°C. The dimer fraction was purified on size exclusion chromatography (Superdex 75) using buffer B. Then it was treated with Yuh1 to remove the proximal D77 residue, purified on size exclusion chromatography, and buffer exchanged into MiliQ water. 3) The dimers obtained from steps 1 and 2 were mixed in the equimolar ratio (50 uM) in the presence of 3 uM E1, 20 uM cdc34, and 3 uM AMSH in reaction buffer A, and incubated overnight at 37°C. *Ub K6/K48/K48 distal: 1)* K6/K48 branched trimer (without the removed D77 from the proximal subunit) was mixed with Ub K48R D77, 1 uM E1, 10 uM UbcH7, 1 uM NleL, 3 uM AMSH. *Ub K6/K48/K63 triple branched were generated in two steps:* 1) 250 uM Ub D77, 500 uM Ub K6R/K48R/K63R, 3uM E1, 10 uM cdc34, 1 uM UbcH13-MmS2 were mixed in buffer A and incubated overnight at 37°C. The resulting trimer was purified using size exclusion chromatography and buffer exchanged into MiliQ water. 2) 40 uM of the trimer obtained in step 1 was mixed with 100 uM Ub K6R/K48R, 3 uM E1, 10 uM UbcH7, 1 uM NleL in buffer A and incubated overnight at 37°C.

All tetramers were purified using size exclusion chromatography (Superdex200) using buffer B (50 mM Tris-HCl pH 7.5, 300 mM NaCl) and buffer exchanged into MiliQ water. Concentrations of all trimers were measured by BCA assay. If applicable, D77 residue was removed from the proximal subunit and size exclusion chromatography was further performed for the second time using Buffer B with the subsequent buffer exchange into MiliQ water.

### Generation of Yuh1-conjugated Ub chains

Autoubiquitination reactions of Yuh1qm5.6 variant were performed at RT in a buffer containing 50 mM HEPES pH 8.0 and 1 mM EDTA. 5 μM of Yuh1qm5.6 was reacted with 50 μM of Ub chains with D77 at the proximal end for 15-30 minutes. The reactions were quenched by incubating with a solution containing 25 mM iodoacetamide (IAA) in the dark for 2-3 hours. Ubn-Yuh1qm5.6 conjugates were purified on size exclusion chromatography (Superdex200) using 50 mM Tris, pH 7.5, 300 mM NaCl, 1 mM DTT, 5% glycerol. Purified Ubn-Yuh1qm5.6 chains were flash frozen and stored in pellets at - 80°C.

### Synthesis of ubiquitinated EGFP species

Ubiquitination reaction was performed by mixing 200 uM EGFP, 200 uM Ub wt, 0.5 uM E1, 10 uM UbcH7, 1 uM NleL in a reaction buffer A (20 mM ATP, 10 mM MgCl2, 40 mM Tris-HCl pH 7.5, 50 mM NaCl, and 0.6 mM DTT) and incubation overnight at 37°C. The reaction mixture was purified 1) by anion exchange chromatography using Buffer A 20 mM Tris pH 8.0, 1 mM DTT and Buffer B 20 mM Tris pH 8.0, 1M NaCl, 1 mM DTT, to remove free Ub chains; 2) by size exclusion chromatography (Superdex200) using Buffer C 50 mM Tris pH 7.5, 150 mM NaCl, 1 mM DTT, 5% glycerol. Purified Ub_n_-GFP chains were flash frozen and stored in pellets at −80°C.

### Asp-N digestion of ubiquitinated substrates

10-20 ug of UbnYuh1qm5.6 or Ub_n_GFP were digested using Asp-N protease with 1:20 protease:substrate mass ratio in reaction buffer A (50 mM ammonium bicarbonate, pH 8.0). For in-gel cleavage efficiency detection the reactions were quenched with SDS-loading dye; for downstream MS experiments − with 0.1% formic acid.

### Top-down analysis of free Ub chains

Prior to MS analysis free Ub chains were additionally desalted using Zeba Spin Desalting Columns (7 MWCO, Thermo Scientific, San Jose, CA) and diluted to a final concentration of 5 uM in 50/49/1 (v/v/v) mixture of acetonitrile/water/formic acid. Intact chains were infused at a rate of 0.25 uL/min using Nano Easy-Spray Emitter (Thermo Scientific, San Jose, CA) and EASY-nLC 1000 System (Thermo Fisher Scientific, San Jose, CA) directly into an Orbitrap Fusion mass spectrometer (Thermo Fisher Scientific, San Jose, CA). All MS^1^ and MS^2^ spectra were acquired at a resolving power of 240k at m/z=200 in intact protein mode with 5 mtorr ion routing multipole pressure. Isolated ions were fragmented with 3 ms electron transfer dissociation supplemented with 10% higher-energy collision dissociation (EThcD), and 10% in in-source activation. Signal accumulation was performed on average for 5-10 minutes. The data quality was assessed using Xcalibur Qual Browser (Thermo Scientific, San Jose, CA).

### Top-down analysis of ubiquitinated substrates

The analysis was performed on LC-MS scale using an Ultimate 3000 ultra-high performance liquid chromatography (Thermo Scientific, San Jose, CA) interfaced to an Orbitrap Fusion mass spectrometer. For the analysis without additional desalting steps ∼1-2 ug (for MS^1^ level) or ∼5 ug (for MS^2^ level) of Ub conjugate(s) was injected onto ZORBAX StableBond 300 C3 column (4.6 x 50 mm, 3.5 µm, Agilent Technologies, Santa Clara, CA) and separated at 0.2 mL/min over 50 minutes using the following gradient: 0-5 min 5% Buffer B, 5-35 minutes 5-40% Buffer B, 35-40 minutes 40-95% Buffer B, 40-42 minutes 95% Buffer B, 42-44 minutes 95-5% Buffer B, 44-50 minutes 5% Buffer B. All MS^1^ and MS^2^ spectra were acquired at a resolving power of 240k at m/z=200 in intact protein mode with 5 mtorr ion routing multipole pressure. MS^2^ analysis was performed in a targeted MS^2^ mode with a precursor list of ions. Isolated ions were fragmented with 3 ms electron transfer dissociation supplemented with 10% higher-energy collision dissociation (EThcD) and 10% in-source activation. The data quality was assessed using Xcalibur Qual Browser (Thermo Scientific, San Jose, CA).

## Supporting information

The Supporting Information is available free of charge.

## ASSOCIATED CONTENT

The Supporting Information is available free of charge. Supplementary Fig.1 shows a machine -readable representation of Ub chains; Supplementary Fig.2 demonstrates how to input information into UbqTop software; Supplementary Fig.3 shows examples of .txt files used to input the sequences of Ub conjugate subunits and the list of the analyzed Ub conjugates; Supplementary Fig.4 shows coverage maps for the characteristic fragment series of Ub dimers; Supplementary Fig.5 shows scoring plots for Ub dimers; Supplementary Fig.6 shows fragment annotation and coverage intervals for ck(Ub)+2Ub fragment series obtained for Ub trimers; Supplementary Fig.7 shows coverage maps for the characteristic fragment series of Ub trimers; Supplementary Fig.8 scoring plots for Ub trimers; Supplementary Fig.9 shows coverage maps for the characteristic fragment series of Ub tetramers; Supplementary Fig.10 shows scoring plots for Ub tetramers; Supplementary Fig.11 demonstrates the Ub chain-modified site correlation problem; Supplementary Fig.12 shows the results of Asp-N digestion of low molecular weight Ub chains; Supplementary Fig.13 shows coverage maps for the characteristic fragment series of Yuh1-Ub2 isomers; Supplementary Fig.14 shows coverage maps for the characteristic fragment series of Yuh1-Ub3 isomers; Supplementary Fig.15 shows coverage maps for the characteristic fragment series for three different GFP fragments modified with two Ub subunits; Supplementary Fig.16 shows TIC for the detected tryptic Ub-GG-peptides. Supplementary Table 1 shows theoretical fragment series and fragment series intervals for Ub dimers; Supplementary Table 2 shows theoretical fragment series and fragment series intervals for Ub trimers; Supplementary Table 3 shows theoretical fragment series and fragment series intervals for Ub tetramers; Supplementary Table 4 shows theoretical fragment series and fragment series intervals for Yuh1qm5.6 modified with Ub dimers; Supplementary Table 5 shows theoretical fragment series and fragment series intervals for Yuh1qm5.6 modified with Ub trimers; Supplementary Table 6 shows characteristic fragment series and fragment series intervals for different EGFP fragments modified with Ub dimers; Supplementary Table 7 shows the results of bottom-up analysis of chain linkage composition for Ubn-GFP mixture; Supplementary Table 8 shows the results of bottom-up analysis of ubiquitination sites for Ubn-GFP mixture.

## CODE AVAILABILITY

All relevant code can be found at https://github.com/dg-ivanov/UbqTop.

## AUTHOR CONTRIBUTIONS

E.R.S. conceived the project, conceptualization, funding acquisition, and supervision. E.S. and J.Z.: generated Ub conjugates, performed MS analysis. D.I., E.S., and L.C.: developed computational algorithms, performed data analyses. E.S., D.I., and E.R.S. wrote the manuscript.

## ACKNOWLEDGEMENTS

This work was supported by research grant R35GM149532 from the National Institutes of Health (NIH). The authors gratefully acknowledge the UMass Amherst Institute of Applied Life Sciences Mass Spectrometry Core Facility (RRID:SCR_019063) for their support and assistance in this work and the equipment funding provided by the Massachusetts Life Science Center. We thank UMass Amherst Institute of Applied Life Sciences Mass Spectrometry Core Facility director Dr. Stephen Eyles for the training and assistance. The data described herein were acquired on an Orbitrap Fusion mass spectrometer funded by NIH grant 1S10OD010645-01A1.

## REFERENCES

1 Mukhopadhyay D, Riezman H. Proteasome-Independent Functions of Ubiquitin in Endocytosis and Signaling. Science (80- ) 2007;315:201–5. 10.1126/science.1127085.

2 Vucic D, Dixit VM, Wertz IE. Ubiquitylation in apoptosis: a post-translational modification at the edge of life and death. Nat Rev Mol Cell Biol 2011;12:439–52. 10.1038/nrm3143.

3 Grabbe C, Husnjak K, Dikic I. The spatial and temporal organization of ubiquitin networks. Nat Rev Mol Cell Biol 2011;12:295–307. 10.1038/nrm3099.

4 Jackson SP, Durocher D. Regulation of DNA Damage Responses by Ubiquitin and SUMO. Mol Cell 2013;49:795–807. 10.1016/j.molcel.2013.01.017.

5 Pickart CM. Mechanisms Underlying Ubiquitination. Annu Rev Biochem 2001;70:503–33. 10.1146/annurev.biochem.70.1.503.

6 Swatek KN, Komander D. Ubiquitin modifications. Cell Res 2016;26:399–422. 10.1038/cr.2016.39.

7 Deol KK, Lorenz S, Strieter ER. Enzymatic Logic of Ubiquitin Chain Assembly. Front Physiol 2019;10:1–14. 10.3389/fphys.2019.00835.

8 Akutsu M, Dikic I, Bremm A. Ubiquitin chain diversity at a glance. J Cell Sci 2016;129:875–80. 10.1242/jcs.183954.

9 Hospenthal MK, Freund SMV, Komander D. Assembly, analysis and architecture of atypical ubiquitin chains. Nat Struct Mol Biol 2013. 10.1038/nsmb.2547.

10 Komander D, Rape M. The Ubiquitin Code. Annu Rev Biochem 2012;81:203–29. 10.1146/annurev-biochem-060310-170328.

11 Damgaard RB. The ubiquitin system: from cell signalling to disease biology and new therapeutic opportunities. Cell Death Differ 2021;28:423–6. 10.1038/s41418-020-00703-w.

12 Tracz M, Bialek W. Beyond K48 and K63: non-canonical protein ubiquitination. Cell Mol Biol Lett 2021;26:1. 10.1186/s11658-020-00245-6.

13 Matsumoto ML, Wickliffe KE, Dong KC, Yu C, Bosanac I, Bustos D, et al. K11-Linked Polyubiquitination in Cell Cycle Control Revealed by a K11 Linkage-Specific Antibody. Mol Cell 2010;39:477–84. 10.1016/j.molcel.2010.07.001.

14 Meyer H-J, Rape M. Enhanced Protein Degradation by Branched Ubiquitin Chains. Cell 2014;157:910–21. 10.1016/j.cell.2014.03.037.

15 Emmerich CH, Ordureau A, Strickson S, Arthur JSC, Pedrioli PGA, Komander D, et al. Activation of the canonical IKK complex by K63/M1-linked hybrid ubiquitin chains. Proc Natl Acad Sci 2013;110:15247–52. 10.1073/pnas.1314715110.

16 Yu Y, Zheng Q, Erramilli SK, Pan M, Park S, Xie Y, et al. K29-linked ubiquitin signaling regulates proteotoxic stress response and cell cycle. Nat Chem Biol 2021;17:896–905. 10.1038/s41589-021-00823-5.

17 Pan M-R, Peng G, Hung W-C, Lin S-Y. Monoubiquitination of H2AX Protein Regulates DNA Damage Response Signaling. J Biol Chem 2011;286:28599–607. 10.1074/jbc.M111.256297.

18 Madiraju C, Novack JP, Reed JC, Matsuzawa S. K63 ubiquitination in immune signaling. Trends Immunol 2022;43:148–62. 10.1016/j.it.2021.12.005.

19 Liu S, Chen ZJ. Expanding role of ubiquitination in NF-κB signaling. Cell Res 2011;21:6–21. 10.1038/cr.2010.170.

20 Fiil BK, Gyrd-Hansen M. The Met1-linked ubiquitin machinery in inflammation and infection. Cell Death Differ 2021;28:557–69. 10.1038/s41418-020-00702-x.

21 Deol KK, Strieter ER. The ubiquitin proteoform problem. Curr Opin Chem Biol 2021;63:95–104. 10.1016/j.cbpa.2021.02.015.

22 Ohtake F. Mass Spectrometry Technologies for Deciphering the Ubiquitin Code. Trends Biochem Sci 2020;45:820–1. 10.1016/j.tibs.2020.04.008.

23 Peng J, Schwartz D, Elias JE, Thoreen CC, Cheng D, Marsischky G, et al. A proteomics approach to understanding protein ubiquitination. Nat Biotechnol 2003;21:921–6. 10.1038/nbt849.

24 Sahu I, Zhu H, Buhrlage SJ, Marto JA. Proteomic approaches to study ubiquitinomics. Biochim Biophys Acta - Gene Regul Mech 2023;1866:194940. 10.1016/j.bbagrm.2023.194940.

25 Hospenthal MK, Freund SM V, Komander D. Assembly, analysis and architecture of atypical ubiquitin chains. Nat Struct Mol Biol 2013;20:555–65. 10.1038/nsmb.2547.

26 Hu Z, Li H, Wang X, Ullah K, Xu G. Proteomic approaches for the profiling of ubiquitylation events and their applications in drug discovery. J Proteomics 2021;231:103996. 10.1016/j.jprot.2020.103996.

27 Valkevich EM, Sanchez NA, Ge Y, Strieter ER. Middle-Down Mass Spectrometry Enables Characterization of Branched Ubiquitin Chains. Biochemistry 2014;53:4979–89. 10.1021/bi5006305.

28 Crowe SO, Rana ASJB, Deol KK, Ge Y, Strieter ER. Ubiquitin Chain Enrichment Middle-Down Mass Spectrometry Enables Characterization of Branched Ubiquitin Chains in Cellulo. Anal Chem 2017;89:4428–34. 10.1021/acs.analchem.6b03675.

29 Xu P, Peng J. Characterization of Polyubiquitin Chain Structure by Middle-down Mass Spectrometry. Anal Chem 2008;80:3438–44. 10.1021/ac800016w.

30 Rana ASJB, Ge Y, Strieter ER. Ubiquitin Chain Enrichment Middle-Down Mass Spectrometry (UbiChEM-MS) Reveals Cell-Cycle Dependent Formation of Lys11/Lys48 Branched Ubiquitin Chains. J Proteome Res 2017;16:3363–9. 10.1021/acs.jproteome.7b00381.

31 Finley D. Recognition and Processing of Ubiquitin-Protein Conjugates by the Proteasome. Annu Rev Biochem 2009;78:477–513. 10.1146/annurev.biochem.78.081507.101607.

32 Winget JM, Mayor T. The Diversity of Ubiquitin Recognition: Hot Spots and Varied Specificity. Mol Cell 2010;38:627–35. 10.1016/j.molcel.2010.05.003.

33 Ikeda F, Crosetto N, Dikic I. What Determines the Specificity and Outcomes of Ubiquitin Signaling? Cell 2010;143:677–81. 10.1016/j.cell.2010.10.026.

34 Lee AE, Geis-Asteggiante L, Dixon EK, Miller M, Wang Y, Fushman D, et al. Preparing to read the ubiquitin code: top-down analysis of unanchored ubiquitin tetramers. J Mass Spectrom 2016;51:629–37. 10.1002/jms.3787.

35 Chen D, Gomes F, Abeykoon D, Lemma B, Wang Y, Fushman D, et al. Top-Down Analysis of Branched Proteins Using Mass Spectrometry. Anal Chem 2018;90:4032–8. 10.1021/acs.analchem.7b05234.

36 Cannon JR, Martinez-Fonts K, Robotham SA, Matouschek A, Brodbelt JS. Top-Down 193-nm Ultraviolet Photodissociation Mass Spectrometry for Simultaneous Determination of Polyubiquitin Chain Length and Topology. Anal Chem 2015;87:1812–20. 10.1021/ac5038363.

37 Geis-Asteggiante L, Lee AE, Fenselau C. Analysis of the topology of ubiquitin chains. Methods Enzymol., vol. 626. 1st ed. Elsevier Inc.; 2019. p. 323–46.

38 Melby JA, Roberts DS, Larson EJ, Brown KA, Bayne EF, Jin S, et al. Novel Strategies to Address the Challenges in Top-Down Proteomics. J Am Soc Mass Spectrom 2021;32:1278–94. 10.1021/jasms.1c00099.

39 Donald E. Knuth. The art of computer programming, volume 1 (3rd ed.): fundamental algorithms. 1997.

40 Tabb DL, Jeong K, Druart K, Gant MS, Brown KA, Nicora C, et al. Comparing Top-Down Proteoform Identification: Deconvolution, PrSM Overlap, and PTM Detection. J Proteome Res 2023;22:2199–217. 10.1021/acs.jproteome.2c00673.

41 Chang LH, Strieter ER. Reprogramming a Deubiquitinase into a Transamidase. ACS Chem Biol 2018;13:2808–18. 10.1021/acschembio.8b00759.

42 Lin DY, Diao J, Zhou D, Chen J. Biochemical and Structural Studies of a HECT-like Ubiquitin Ligase from Escherichia coli O157:H7. J Biol Chem 2011;286:441–9. 10.1074/jbc.M110.167643.

43 Pickart CM, Raasi S. Controlled Synthesis of Polyubiquitin Chains. 2005. p. 21–36.

