## Supplementary material for "Computationally Driven Top-Down Mass Spectrometry of Ubiquitinated Proteins": The Supporting Information is available free of charge.

### SUPPLEMENTARY FIGURES

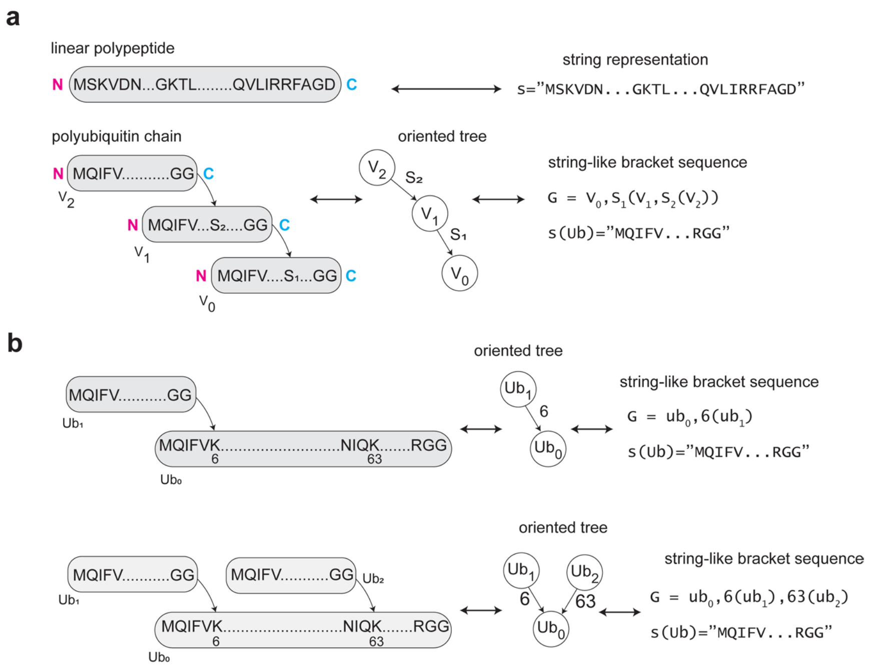


**Supplementary Fig. 1.** Machine-readable representation of standard linear polypeptide and Ub chain. (a) Linear polypeptide can be represented in a "string" format, whereas Ub chain-oriented tree can be represented as a linear bracket sequence. (b) Examples of K6- Ub dimer and K6/K63-branched trimer represented in a bracket sequence format.

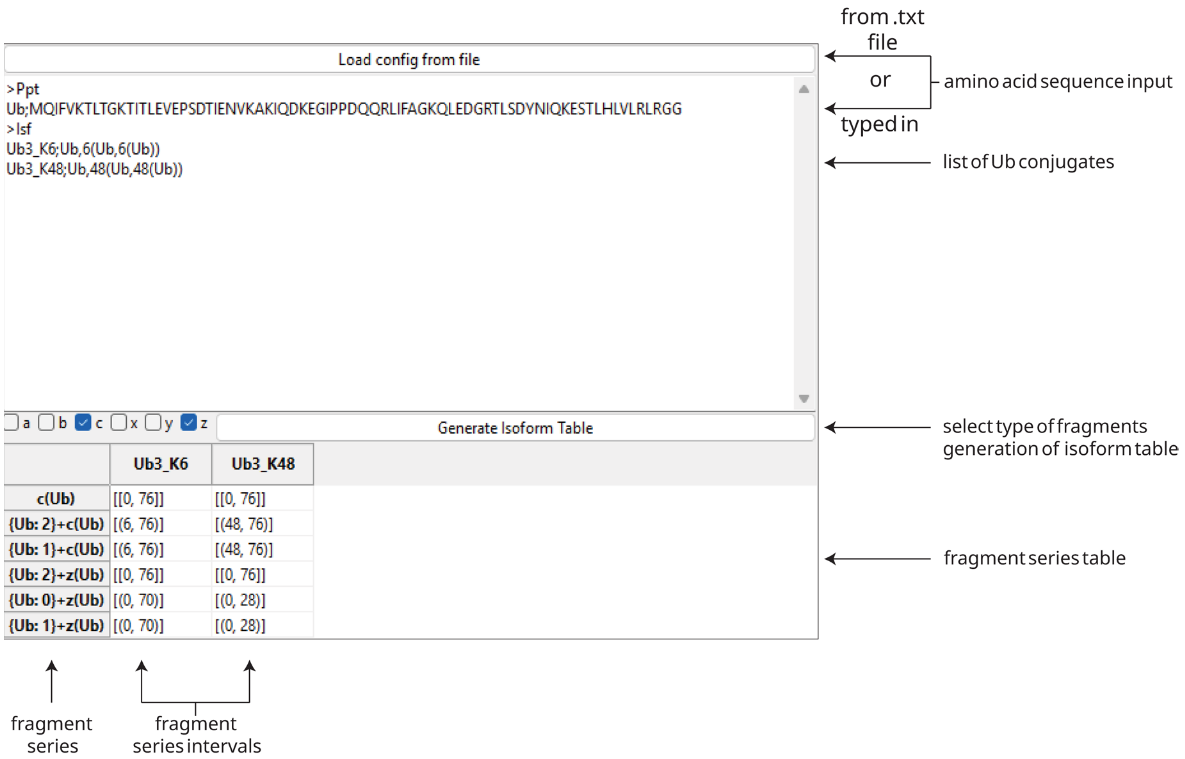


**Supplementary Fig. 2** Entering the information into *UbqTop*: 1) the list of the analyzed Ub conjugates and the sequences of their corresponding subunits can be imported by uploading a .txt. file, or by manually typing in the information into the *UbqTop*. 2) by selecting type of fragmentation, corresponding fragment series and their intervals are generated in a table format.

**a**

```
>Ppt
Ub;MQIFVKTLTGKTITLEVEPSDTIENVKAKIQDKEGIPPDQQRLIFAGKQLEDGRTLSDYNIQKESTLHLVLRLRGG
>Isf
Ub3_K6;Ub,6(Ub,6(Ub))
Ub3_K48;Ub,48(Ub,48(Ub))
Ub3_K63;Ub,63(Ub,63(Ub))
```

**b**

```
>Ppt
Ub;MQIFVKTLTGKTITLEVEPSDTIENVKAKIQDKEGIPPDQQRLIFAGKQLEDGRTLSDYNIQKESTLHLVLRLRGG
Ub_K6R_K48R;MQIFVRTLTGKTITLEVEPSDTIENVKAKIQDKEGIPPDQQRLIFAGRQLEDGRTLSDYNIQKESTLHLVLRLRGG
>Isf
Ub3_K6;Ub,6(Ub,6(Ub))
Ub3_K48;Ub,48(Ub,48(Ub))
Ub3_K6_K48_branched;Ub,6(Ub_K6R_K48R),48(Ub_K6R_K48R)
```

**c**

```
>Ppt
Ub;MQIFVKTLTGKTITLEVEPSDTIENVKAKIQDKEGIPPDQQRLIFAGKQLEDGRTLSDYNIQKESTLHLVLRLRGG
Substrate_Yuh1;DQFVLNVIKENVQTFSTGQSEAPEATAAKKAHYITYVEENGGIFEL
>Isf
Yuh1_site30_Ub2K6;Substrate_Yuh1,30(Ub,6(Ub))
Yuh1_site30_Ub2K48;Substrate_Yuh1,30(Ub,48(Ub))
```

**Supplementary Fig. 3.** Examples of .txt files used to input the sequences of Ub conjugate subunits and the list of the analyzed Ub conjugates. (A) A list of three Ub trimers, consisting of Ub *wild type* only. (B) A list of three Ub trimers, consisting of Ub *wild type* or Ub K6R/K48R double mutant. (C) A list of two ubiquitinated substrates, conjugated to different Ub dimers. As an input sequences of Ub *wild type* and a substrate were added.

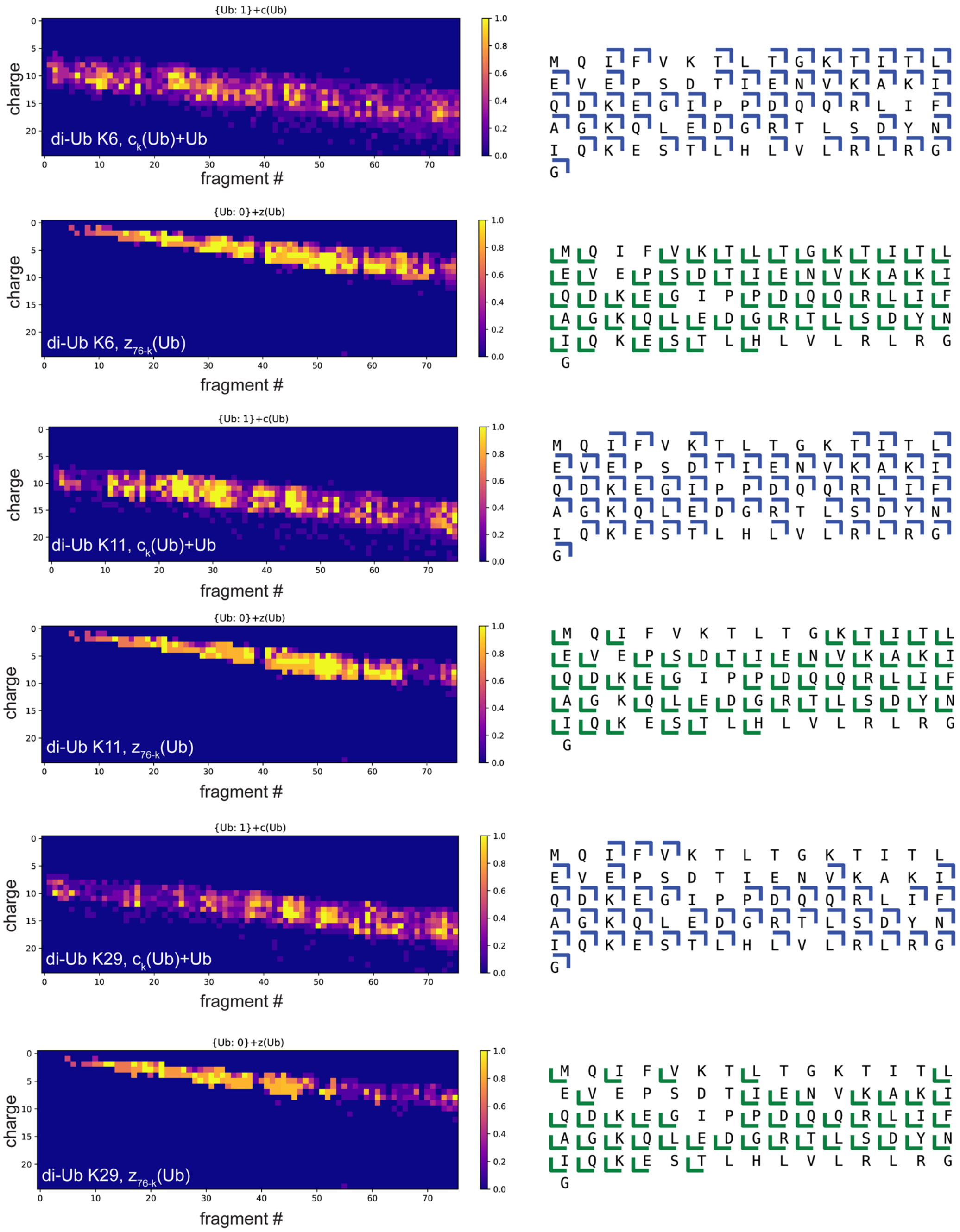


*Continues on the next page, legend is on the next page

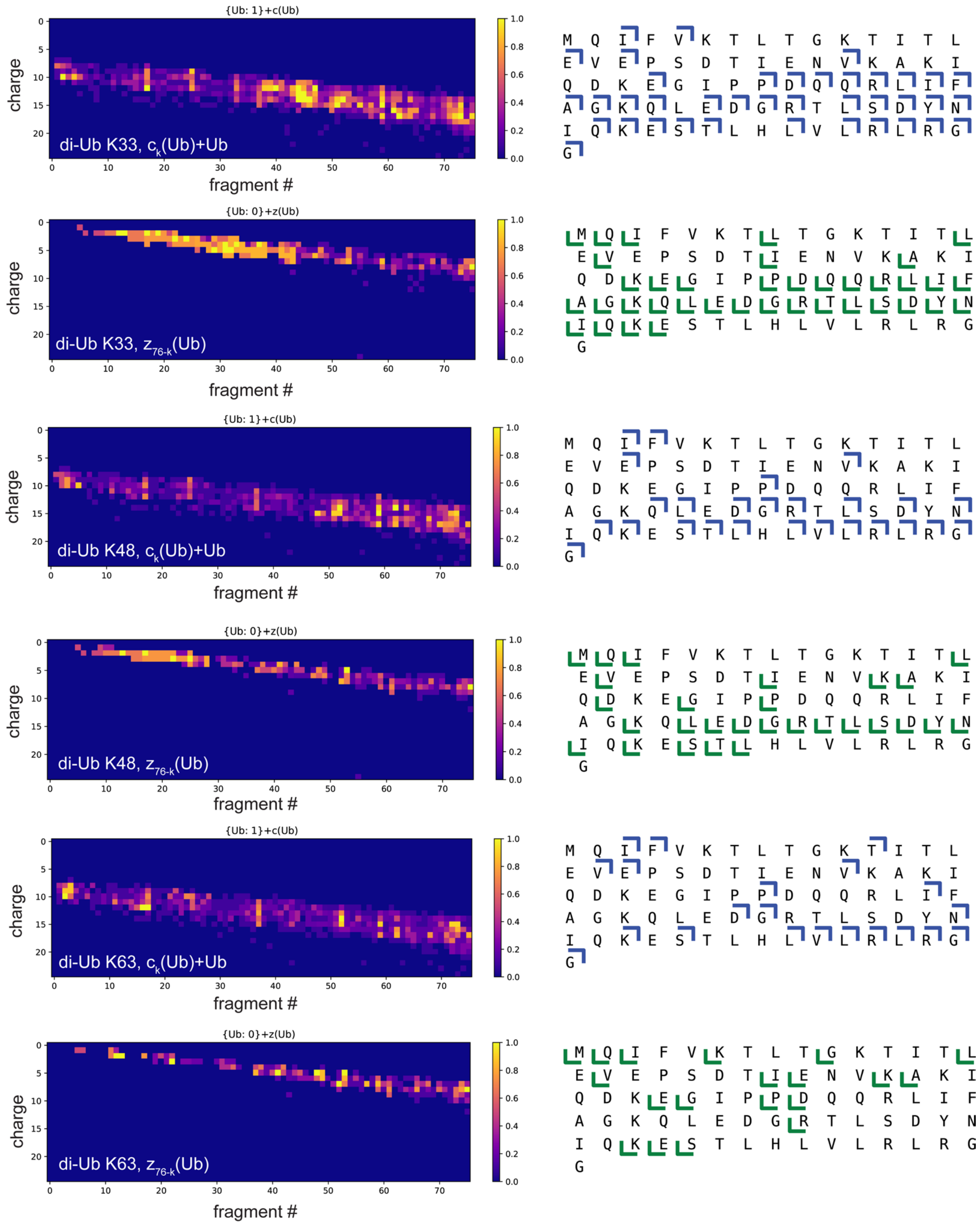


**Supplementary Fig. 4.** Coverage maps for the characteristic fragment series of *Ub dimers*. The left map represents the fragment annotation result. Each colored square is a fragment *k* at a single charge state. The color relates to a fragment quality. Multiple squares positioned vertically represent the same fragment but at different charge states. The right map represents the final coverage interval for the analyzed fragment series obtained after the analysis of all fragments at all detected charge states.

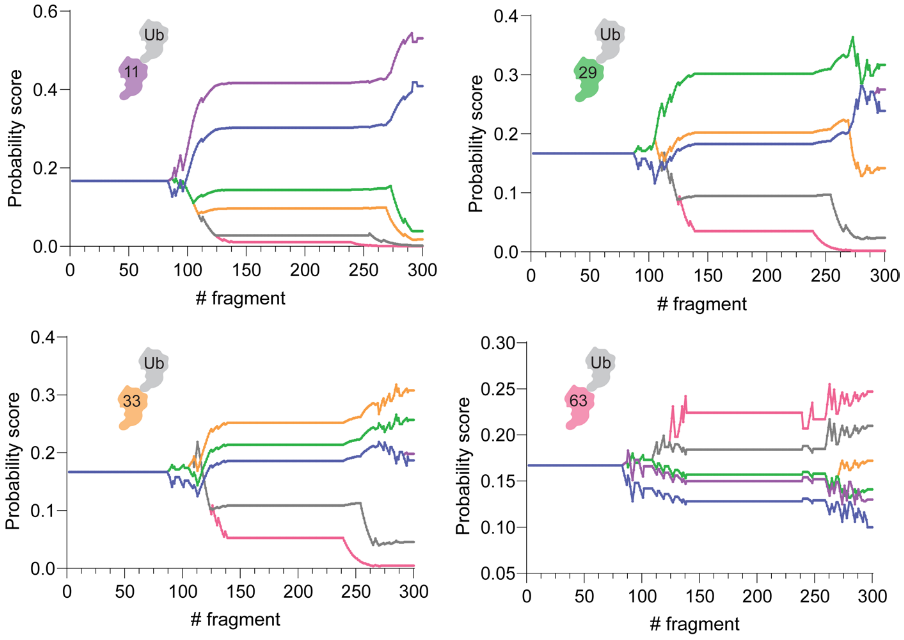


**Supplementary Fig. 5.** Scoring plots for Ub dimers (K11-, K29, K33-, K63).

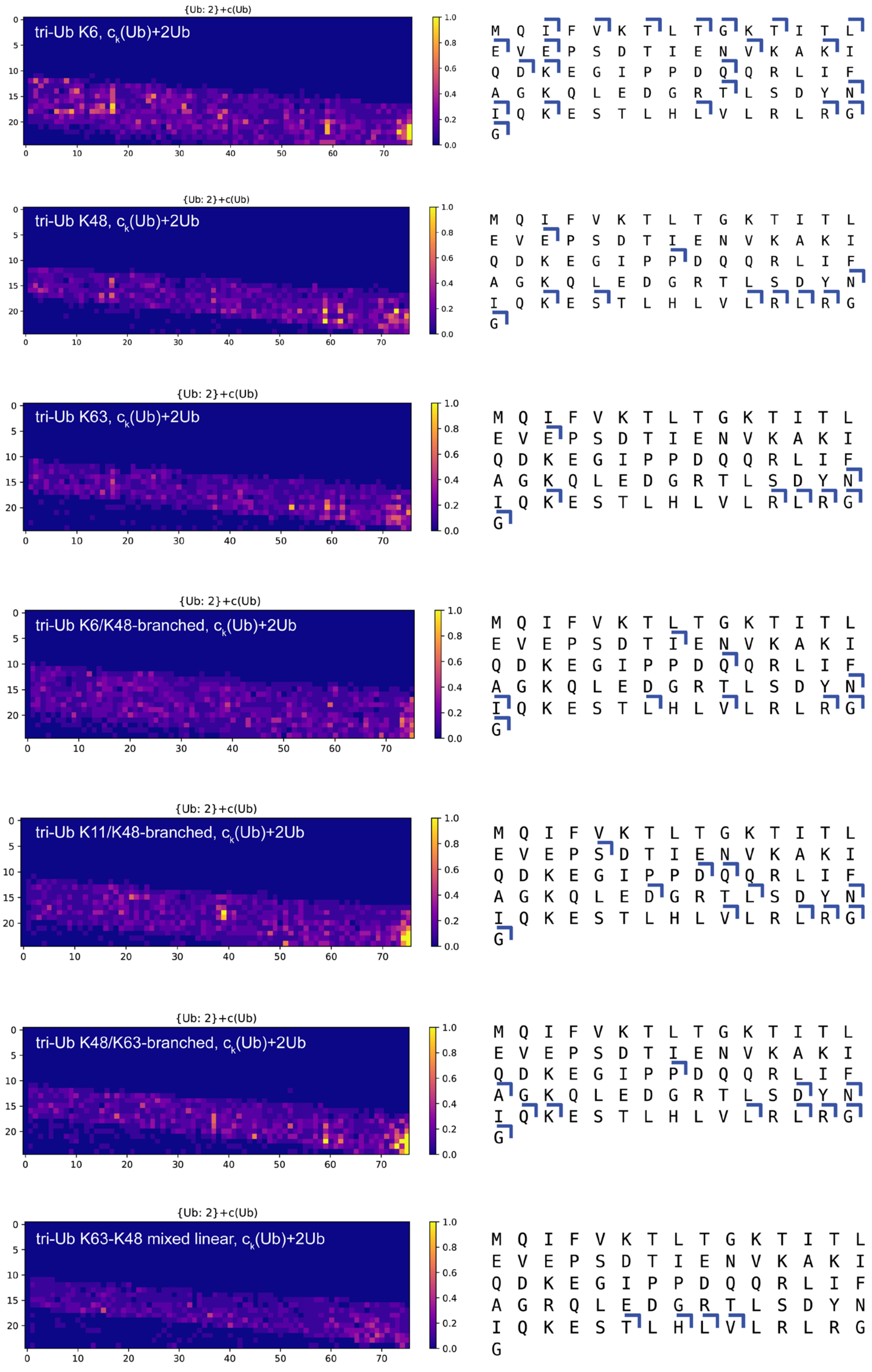


**Supplementary Fig. 6.** Fragment annotation and coverage intervals for $c_k$(Ub)+2Ub fragment series obtained for *Ub trimers*.

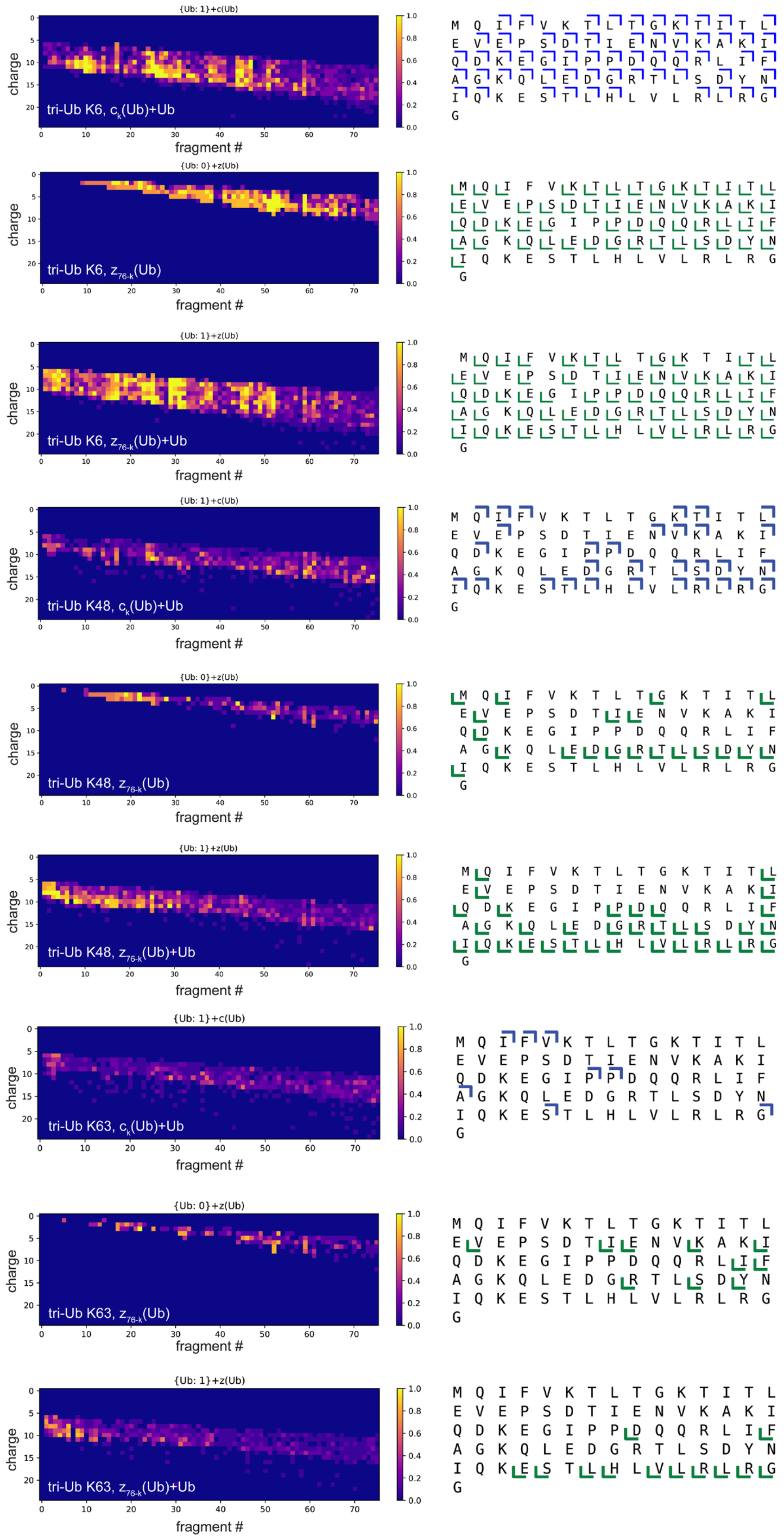


*Continues on the next page

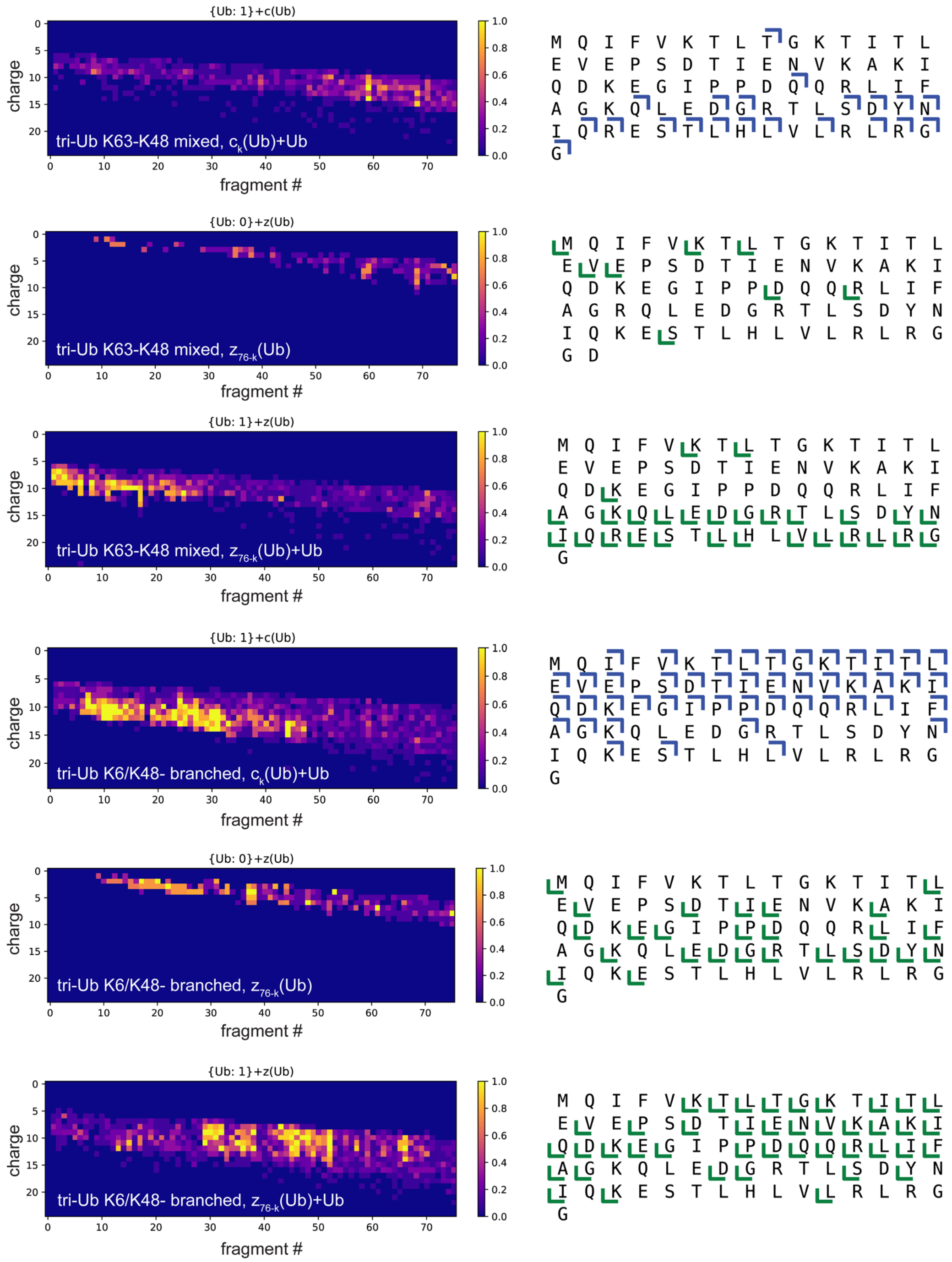


*Continues on the next page, legend is on the next page

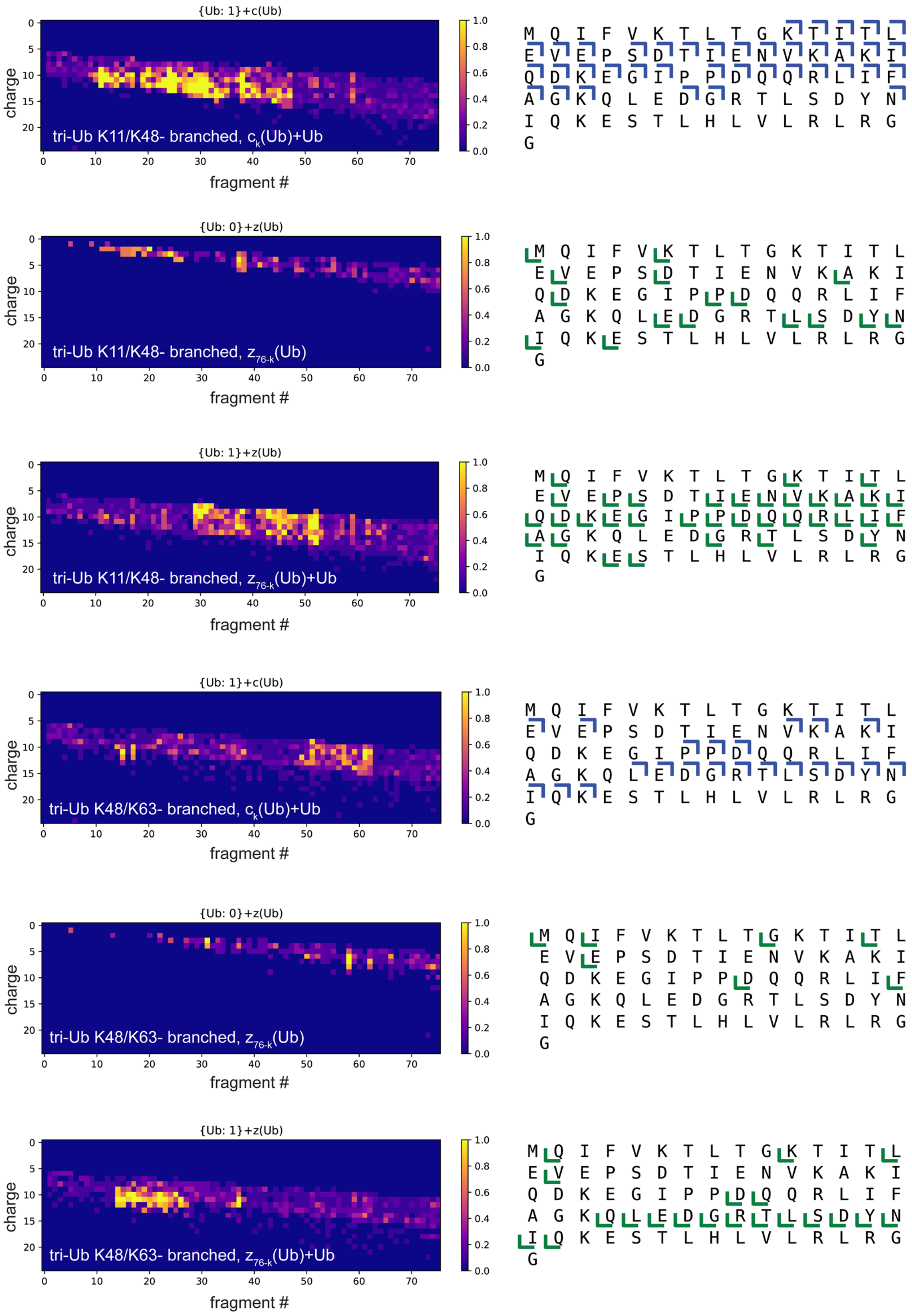


**Supplementary Fig. 7.** Coverage maps for the characteristic fragment series of *Ub trimers*. The left map represents the fragment annotation result. Each colored square is a fragment *k* at a single charge state. The color relates to a fragment quality. Multiple squares positioned vertically represent the same fragment but at different charge states.

The right map represents the final coverage interval for the analyzed fragment series obtained after the analysis of all fragments at the detected charge states.

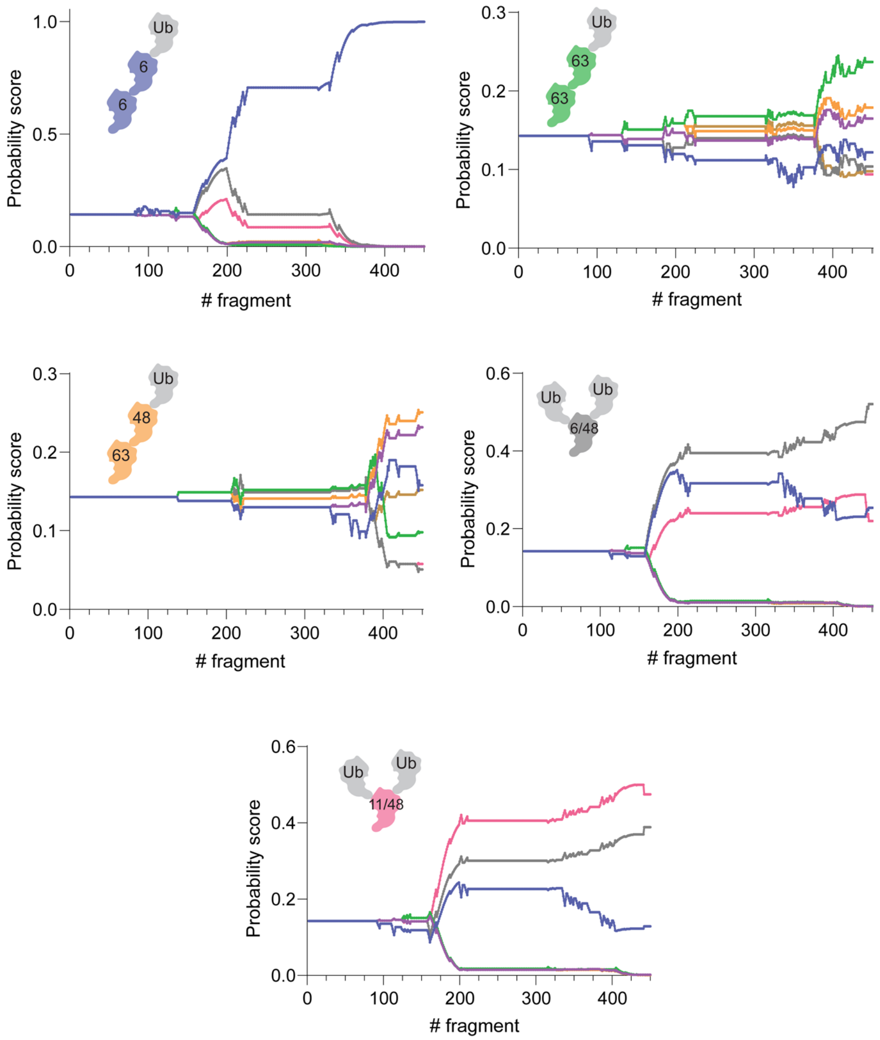


**Supplementary Fig. 8.** Scoring plots for Ub trimers (linear: K6-, K63-, K63-K48 mixed; branched: K6/K48-, K11/K48).

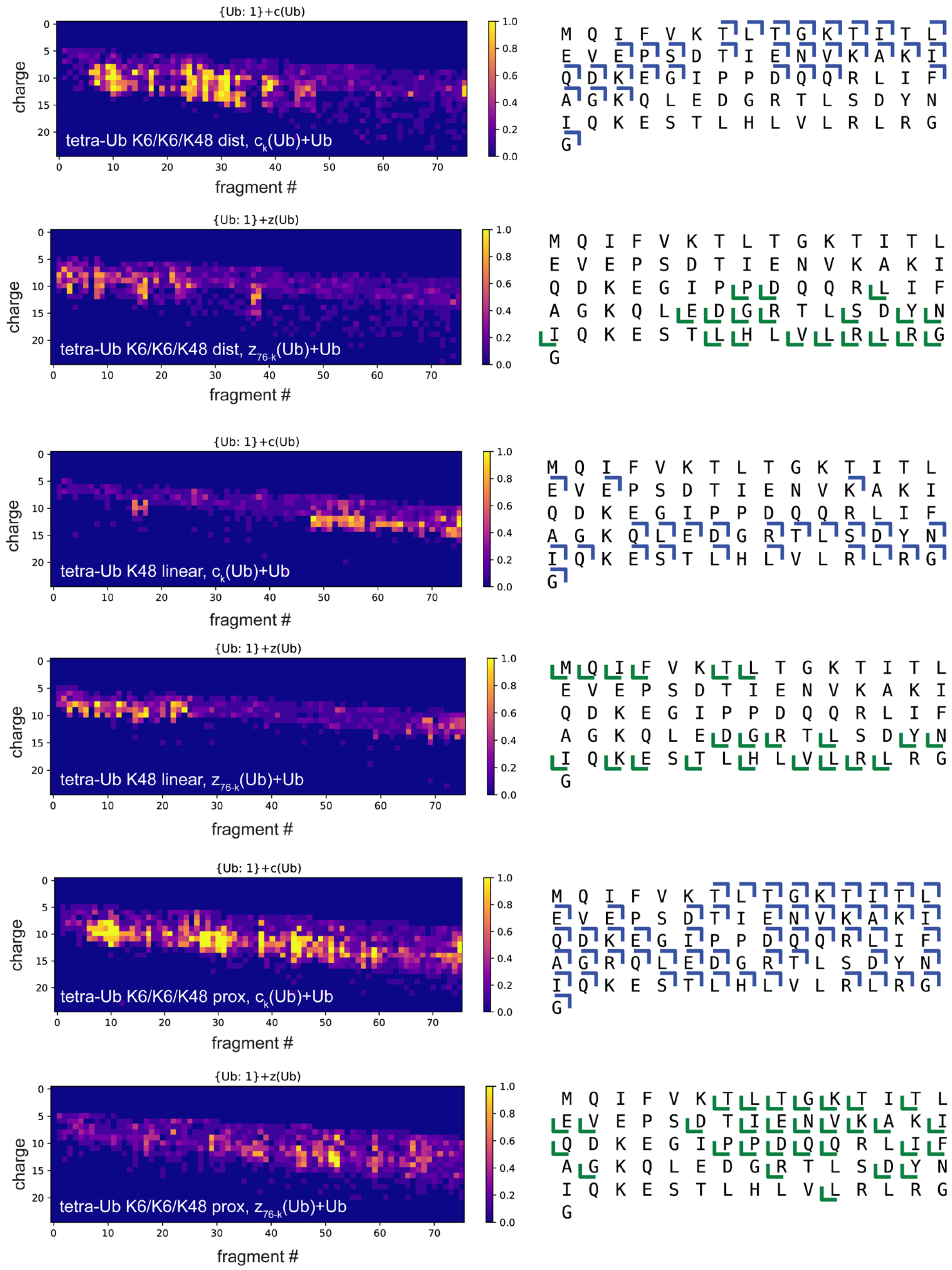


*Continues on the next page, legend is on the next page

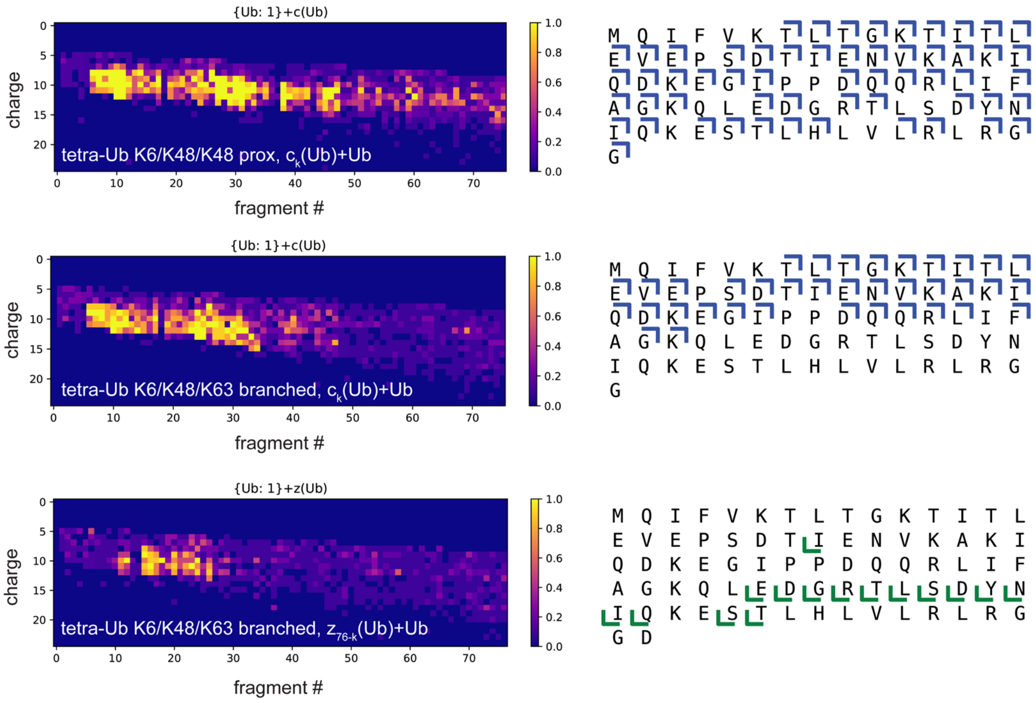


**Supplementary Fig. 9.** Coverage maps for the characteristic fragment series of *Ub tetramers*. The left map represents the fragment annotation result. Each colored square is a fragment k at a single charge state. The color relates to a fragment quality. Multiple squares positioned vertically represent the same fragment but at different charge states. The right map represents the final coverage interval for the analyzed fragment series obtained after the analysis of all fragments at the detected charge states.

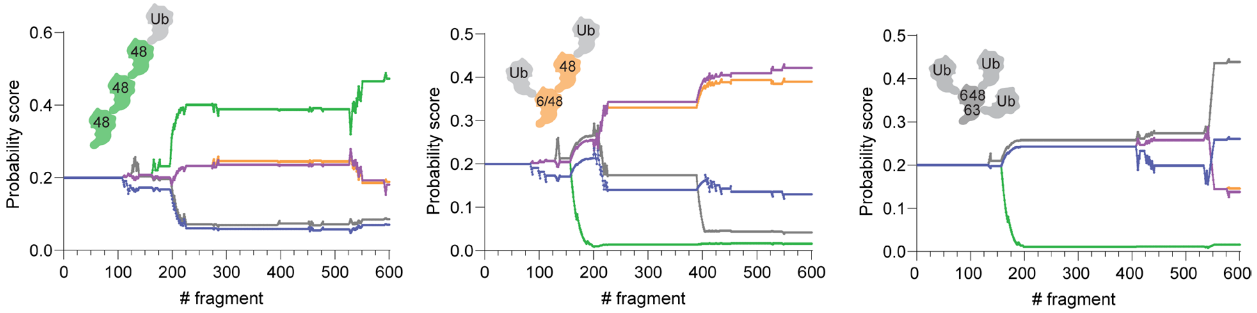


**Supplementary Fig. 10.** Scoring plots for Ub tetramers (linear: K48-; branched: K6/K48/K48 proximal, K6/K48/K63 triple branched).

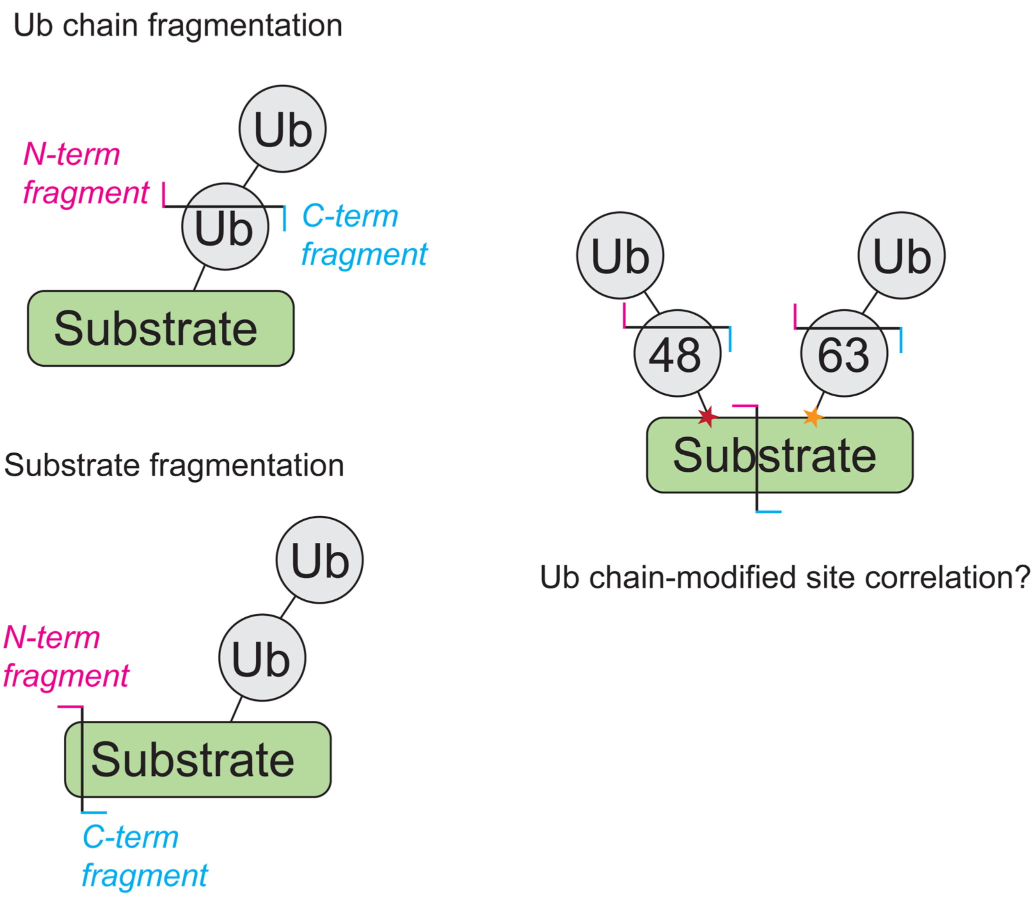


**Supplementary Fig. 11.** With the single cleavage assumption, in case of multiple isomeric chains modifying a substrate the information on the correlation between Ub chain and modified site cannot be obtained at $MS^2$ level.

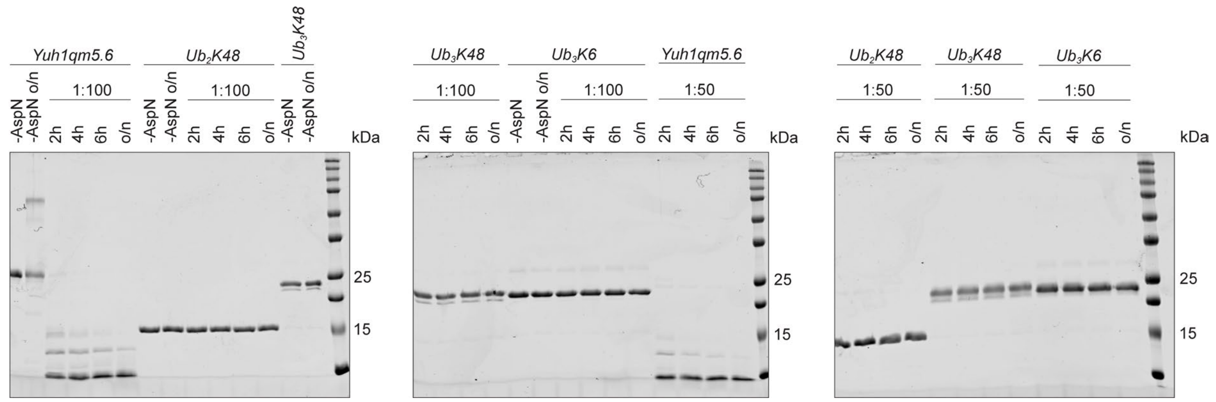


**Supplementary Fig. 12.** Asp-N digestion of low molecular weight (LMW) Ub chains at different protease:substrate ratios (1:100, 1:50) and time points (2 h, 4 h, 6 h, o/n). Yuh1qm5.6 was used as a control to confirm the activity of Asp-N.

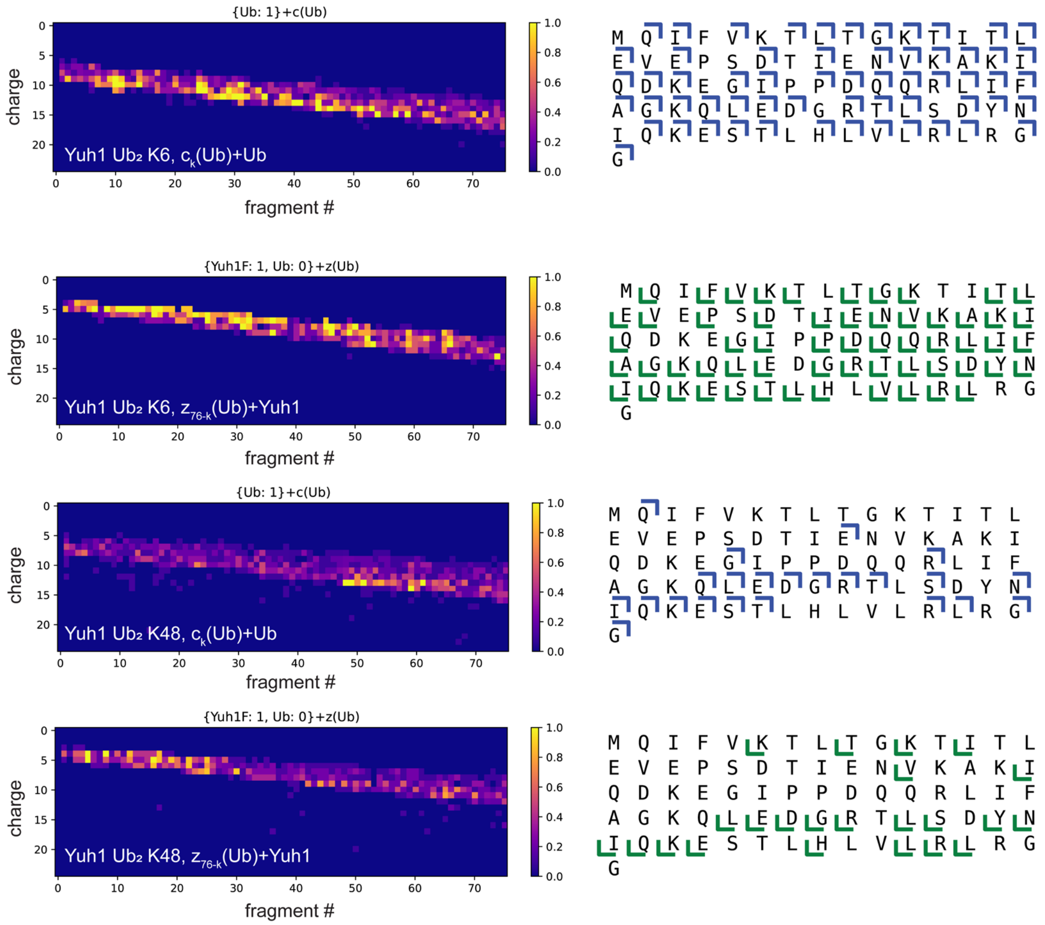


**Supplementary Fig. 13.** Coverage maps for the characteristic fragment series of *Yuh1-Ub2 isomers*. The left map represents the fragment annotation result. Each colored square is a fragment k at a single charge state. The color relates to a fragment quality. Multiple squares positioned vertically represent the same fragment but at different charge states. The right map represents the final coverage interval for the analyzed fragment series obtained after the analysis of all fragments at the detected charge states.

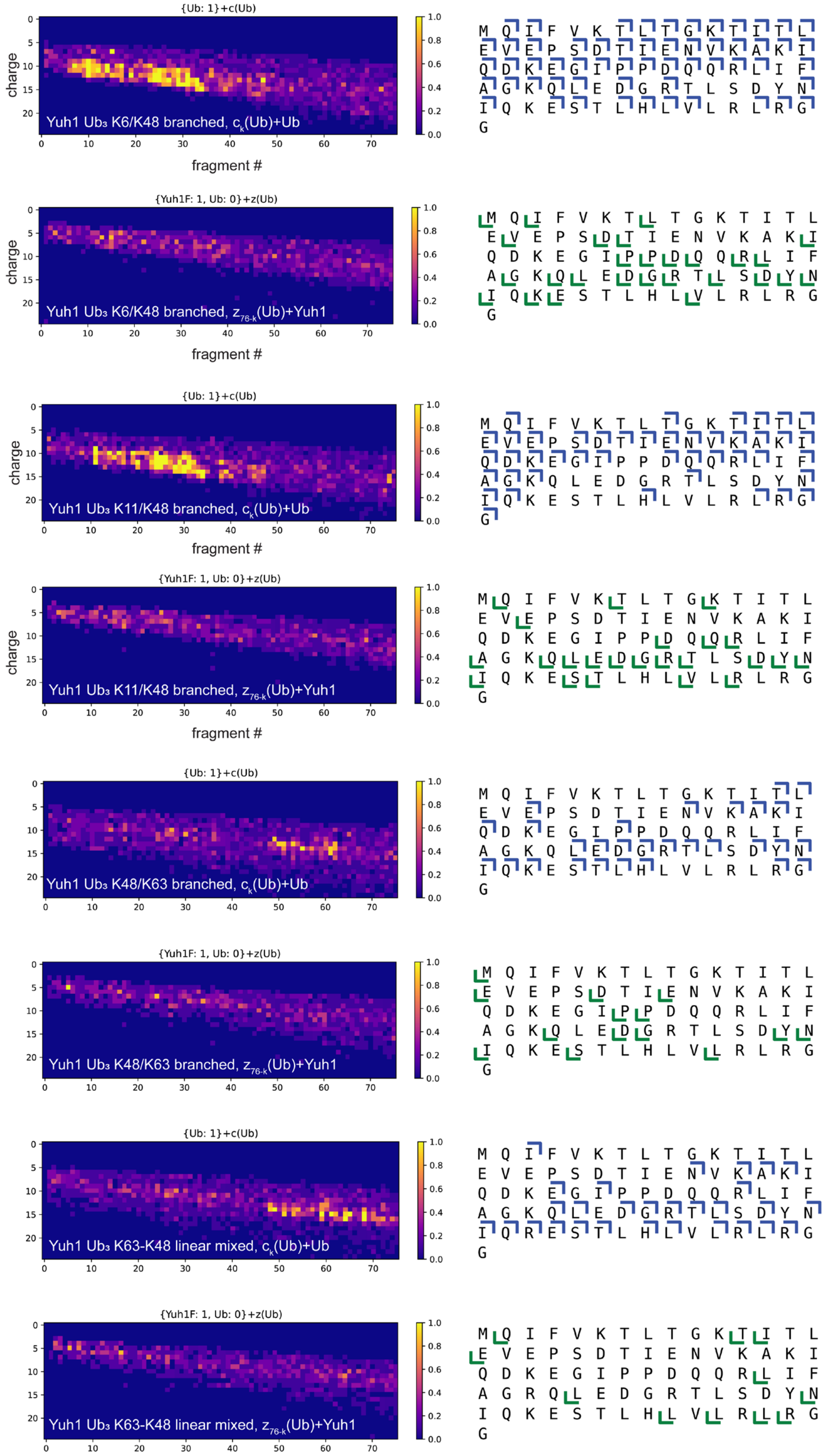


**Supplementary Fig. 14.** Coverage maps for the characteristic fragment series of *Yuh1-Ub3 isomers*. The left map represents the fragment annotation result. Each colored square is a fragment k at a single charge state. The color relates to a fragment quality.

Multiple squares positioned vertically represent the same fragment but at different charge states. The right map represents the final coverage interval for the analyzed fragment series obtained after the analysis of all fragments at the detected charge states.

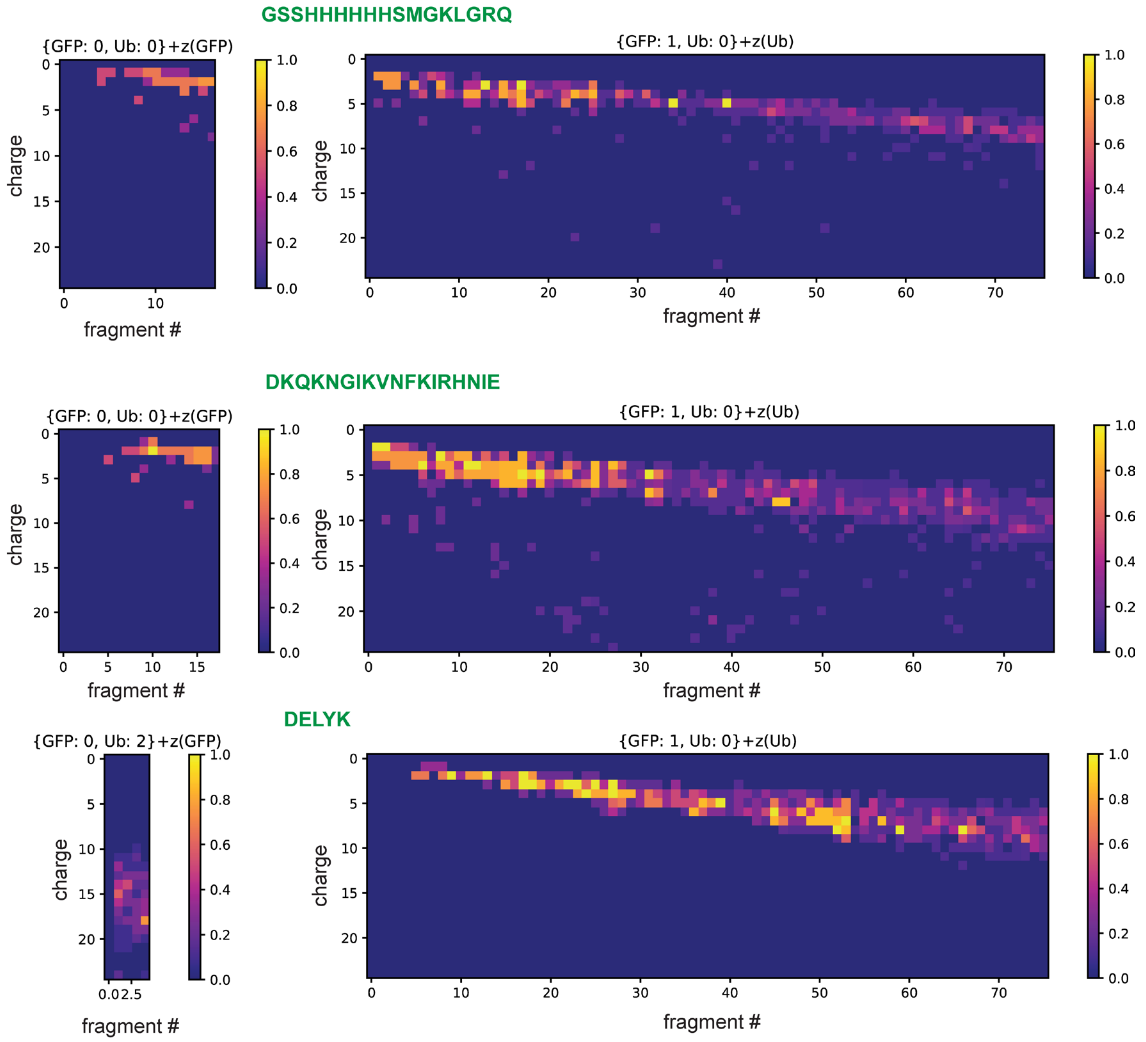


**Supplementary Fig. 15.** Coverage maps for the characteristic fragment series for three different *GFP fragments modified with two Ub subunits*. The left map represents the fragment annotation result. Each colored square is a fragment k at a single charge state. The color relates to a fragment quality. Multiple squares positioned vertically represent the same fragment but at different charge states. The right map represents the final coverage interval for the analyzed fragment series obtained after the analysis of all fragments at the detected charge states.

*K6-GG*

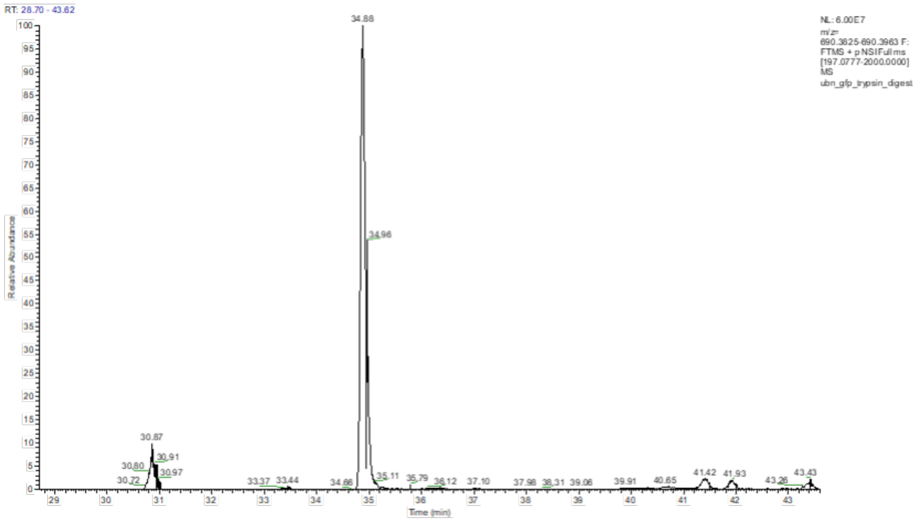


*K48-GG*

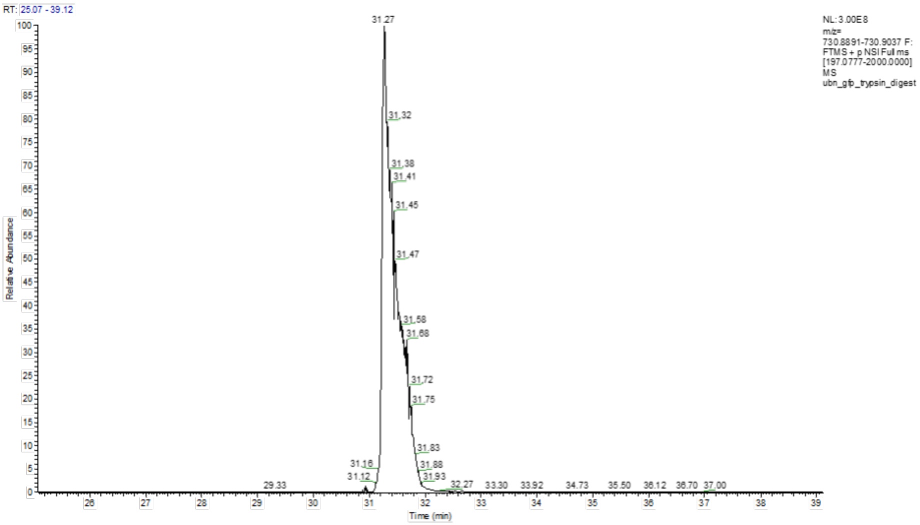


**Supplementary Fig. 16.** TIC for the detected tryptic Ub-GG-peptides.

### SUPPLEMENTARY TABLES

**Supplementary Table 1.** Theoretical fragment series and fragment series intervals for *Ub dimers* (K6-, K11-, K29-, K33-, K48-, K63-). The highlighted series obtain different fragment series intervals and can serve as characteristic series to distinguish between isomers.

| | K6- | K11- | K29- | K33- | K48- | K63- |
|---|---|---|---|---|---|---|
| $C_k$(Ub) | [1, 76) | [1, 76) | [1, 76) | [1, 76) | [1, 76) | [1, 76) |
| ***$C_k$(Ub)+Ub*** | ***[6, 76)*** | ***[11, 76)*** | ***[29, 76)*** | ***[33, 76)*** | ***[48, 76)*** | ***[63, 76)*** |
| ***$Z_{76-k}$*** | ***[1, 70)*** | ***[1, 65)*** | ***[1, 47)*** | ***[1, 43)*** | ***[1, 28)*** | ***[1, 13)*** |
| $Z_{76-k}$(Ub)+Ub | [1, 76) | [1, 76) | [1, 76) | [1, 76) | [1, 76) | [1, 76) |

**Supplementary Table 2.** Theoretical fragment series and fragment series intervals for *Ub trimers* (linear: K6-, K48-, K63-, K63-K48 mixed, branched: K6/K48-, K11/K48-, K48/K63). The highlighted series obtain different fragment series intervals and can serve as characteristic series to distinguish between isomers.

| | K6-linear | K48-linear | K63-linear | K63-K48 mixed | K6/K48-branched | K11/K48-branched | K48/K63-branched |
|---|---|---|---|---|---|---|---|
| $C_k$(Ub) | [1, 76) | [1, 76) | [1, 76) | [1, 76) | [1, 76) | [1, 76) | [1, 76) |
| ***$C_k$(Ub)+2Ub*** | ***[6, 76)*** | ***[48, 76)*** | ***[63, 76)*** | ***[63, 76)*** | ***[48, 76)*** | ***[48, 76)*** | ***[63, 76)*** |
| ***$C_k$(Ub)+Ub*** | ***[6, 76)*** | ***[48, 76)*** | ***[63, 76)*** | ***[48, 76)*** | ***[6, 48)*** | ***[11, 48)*** | ***[48, 63)*** |
| $Z_{76-k}$(Ub)+2Ub | [1, 76) | [1, 76) | [1, 76) | [1, 76) | [1, 76) | [1, 76) | [1, 76) |
| ***$Z_{76-k}$(Ub)*** | ***[1, 70)*** | ***[1, 28)*** | ***[1, 13)*** | ***[1, 13)*** | ***[1, 28)*** | ***[1, 28)*** | ***[1, 13)*** |
| ***$Z_{76-k}$(Ub)+Ub*** | ***[1, 70)*** | ***[1, 28)*** | ***[1, 13)*** | ***[1, 28)*** | ***[28, 70)*** | ***[28, 65)*** | ***[13, 28)*** |

**Supplementary Table 3.** Theoretical fragment series and fragment series intervals for Ub tetramers (K6/K6/K48 distal, K6/K6/K48 proximal, K48- linear, K6/K48/K48 proximal, K6/K48/K63- triple branched). The highlighted series obtain different fragment series intervals and can serve as characteristic series to distinguish between isomers.

| | K6/K6/K48 distal | K6/K6/K48 proximal | K48 linear | K6/K48/K48 proximal | K6/K48/K63 triple branched |
|---|---|---|---|---|---|
| $C_k$(Ub) | [1, 76) | [1, 76) | [1, 76) | [1, 76) | [1, 76) |
| ***$C_k$(Ub)+3Ub*** | ***[6, 76)*** | ***[48, 76)*** | ***[48, 76)*** | ***[48, 76)*** | ***[63, 76)*** |
| ***$C_k$(Ub)+Ub*** | ***[6, 48)*** | ***[6, 76)*** | ***[48, 76)*** | ***[6, 76)*** | ***[6, 48)*** |
| ***Ck(Ub)+2Ub*** | ***[6, 48)*** | ***[6, 48)*** | ***[48, 76)*** | ***-*** | ***[48, 63)*** |
| $Z_{76\text{-}k}$(Ub)+3Ub | [1, 76) | [1, 76) | [1, 76) | [1, 76) | [1, 76) |
| ***$Z_{76\text{-}k}$(Ub)*** | ***[1, 70)*** | ***[1, 28)*** | ***[1, 28)*** | ***[1, 28)*** | ***[1, 13)*** |
| ***$Z_{76\text{-}k}$(Ub)+2Ub*** | ***[28, 70)*** | ***[1, 70)*** | ***[1, 28)*** | ***[1, 70)*** | ***[28, 70)*** |
| **$Z_{76\text{-}k}$(Ub)+Ub** | **[1, 28)** | **[28, 70)** | **[1, 28)** | **-** | **[13, 28)** |

**Supplementary Table 4.** Theoretical fragment series and fragment series intervals for Yuh1qm5.6 modified with Ub dimers (K6- or K48-) at the lysine residue K164 (in Yuh1qm5.6 Asp-N fragment the K number changed to K30). The highlighted series obtain different fragment series intervals and can serve as characteristic series to distinguish between isomers.

| | Yuh1-K6 $Ub_2$ | Yuh1-K48 $Ub_2$ |
|---|---|---|
| $C_k$(Yuh1) | [1, 30) | [1, 30) |
| $C_k$(Yuh1)+2Ub | [30, 46) | [30, 46) |
| $C_k$(Ub) | [1, 76) | [1, 76) |
| ***$C_k$(Ub)+Ub*** | ***[6, 76)*** | ***[48, 76)*** |
| $C_k$(Yuh1)+2Ub | [16, 46) | [16, 46) |
| $Z_{n-k}$(Yuh1) | [1, 16) | [1, 16) |
| $Z_{76-k}$(Ub)+Yuh1+Ub | [1, 76) | [1, 76) |
| ***$Z_{76-k}$(Ub)+Yuh1*** | ***[1, 70)*** | ***[1, 28)*** |

**Supplementary Table 5.** Theoretical fragment series and fragment series intervals for Yuh1qm5.6 modified with Ub trimers (K6/K48-, K11/K48-, K48/K63- branched or K63-K48 linear mixed) at the lysine residue K164 (in Yuh1qm5.6 Asp-N fragment, the K number changed to K30). The highlighted series obtain different fragment series intervals and can serve as characteristic series to distinguish between isomers.

| | Yuh1-K6/K48 branched $Ub_3$ | Yuh1-K11/K48 branched $Ub_3$ | Yuh1-K48/K63 branched $Ub_3$ | Yuh1-K63-K48 mixed linear $Ub_3$ |
|---|---|---|---|---|
| $C_k$(Yuh1) | [1, 30) | [1, 30) | [1, 30) | [1, 30) |
| $C_k$(Yuh1)+3Ub | [30, 46) | [30, 46) | [30, 46) | [30, 46) |
| $C_k$(Ub) | [1, 76) | [1, 76) | [1, 76) | [1, 76) |
| ***$C_k$(Ub)+Ub*** | ***[6, 48)*** | ***[11, 48)*** | ***[48, 63)*** | ***[48, 76)*** |
| ***$C_k$(Ub)+2Ub*** | ***[48, 76)*** | ***[48, 76)*** | ***[63, 76)*** | ***[63, 76)*** |
| $Z_{n-k}$(Yuh1)+3Ub | [16, 46) | [16, 46) | [16, 46) | [16, 46) |
| $Z_{n-k}$(Yuh1) | [1, 16) | [1, 16) | [1, 16) | [1, 16) |
| $Z_{76-k}$(Ub)+Yuh1+2Ub | [1, 76) | [1, 76) | [1, 76) | [1, 76) |
| ***$Z_{76-k}$(Ub)+Yuh1+Ub*** | ***[28, 70)*** | ***[28, 65)*** | ***[13, 28)*** | ***[1, 28)*** |
| ***$Z_{76-k}$(Ub)+Yuh1*** | ***[1, 28)*** | ***[1, 28)*** | ***[1, 13)*** | ***[1, 13)*** |

**Supplementary Table 6.** Characteristic fragment series and fragment series intervals for different EGFP fragments modified with Ub dimers (K6-, K11-, K48-, K63-).

GFP fragment:
***G****SSHHHHHHSMG***K***LGRQ(D)*

| | GFP[K]-K6 $Ub_2$ | GFP[K]-K11 $Ub_2$ | GFP[K]-K48 $Ub_2$ | GFP[K]-K63 $Ub_2$ | GFP[G]-K6 $Ub_2$ | GFP[G]-K11 $Ub_2$ | GFP[G]-K48 $Ub_2$ | GFP[G]-K63 $Ub_2$ |
|---|---|---|---|---|---|---|---|---|
| $Z_{n-k}$(GFP) | [1, 4) | [1, 4) | [1, 4) | [1, 4) | [1, 16) | [1, 16) | [1, 16) | [1, 16) |
| $Z_{76-k}$(Ub) +GFP frag | [1, 70) | [1, 65) | [1, 28) | [1, 13) | [1, 70) | [1, 65) | [1, 28) | [1, 13) |

GFP fragment :
*D***K***QKNGIKVNFKIRHNIE*

| | GFP-K6 $Ub_2$ | GFP-K11 $Ub_2$ | GFP-K48 $Ub_2$ | GFP-K63 $Ub_2$ |
|---|---|---|---|---|
| $Z_{n-k}$(GFP) | [1, 16) | [1, 16) | [1, 16) | [1, 16) |
| $Z_{76-k}$(Ub)+GFP frag | [1, 70) | [1, 65) | [1, 28) | [1, 13) |

GFP fragment :
*DKQ***K***NGIKVNFKIRHNIE*

| | GFP-K6 $Ub_2$ | GFP-K11 $Ub_2$ | GFP-K48 $Ub_2$ | GFP-K63 $Ub_2$ |
|---|---|---|---|---|
| $Z_{n-k}$(GFP) | [1, 14) | [1, 14) | [1, 14) | [1, 14) |
| $Z_{76-k}$(Ub)+GFP frag | [1, 70) | [1, 65) | [1, 28) | [1, 13) |

GFP fragment :
*DKQKNGI***K***VNFKIRHNIE*

| | GFP-K6 $Ub_2$ | GFP-K11 $Ub_2$ | GFP-K48 $Ub_2$ | GFP-K63 $Ub_2$ |
|---|---|---|---|---|
| $Z_{n-k}$(GFP) | [1, 10) | [1, 10) | [1, 10) | [1, 10) |
| $Z_{76-k}$(Ub)+GFP frag | [1, 70) | [1, 65) | [1, 28) | [1, 13) |

GFP fragment :
*DKQKNGIKVNF***K***IRHNIE*

| | GFP-K6 $Ub_2$ | GFP-K11 $Ub_2$ | GFP-K48 $Ub_2$ | GFP-K63 $Ub_2$ |
|---|---|---|---|---|
| $Z_{n-k}$(GFP) | [1, 6) | [1, 6) | [1, 6) | [1, 6) |
| $Z_{76-k}$(Ub)+GFP frag | [1, 70) | [1, 65) | [1, 28) | [1, 13) |

***During multi-monoubiquitination** $C_k$(Ub)+Ub and $Z_{76-k}$(Ub)+GFP fragment series cannot be produced, thus we excluded multi-monoubiquitination as we detect these fragment series.

GFP fragment : *DELY***K**

| | GFP-K6 $Ub_2$ | GFP-K11 $Ub_2$ | GFP-K48 $Ub_2$ | GFP-K63 $Ub_2$ |
|---|---|---|---|---|
| $Z_{n-k}$(GFP)+2Ub | [1, 5) | [1, 5) | [1, 5) | [1, 5) |
| $Z_{76-k}$(Ub)+GFP frag | [1, 70) | [1, 65) | [1, 28) | [1, 13) |

**Supplementary Table 7.** Bottom-up analysis of chain linkage composition for $Ub_n$-GFP mixture.

| | Peptide | Intensity | m/z, z=2 |
|---|---|---|---|
| **K6-GG** | MQIFVKTLTGKGG | 6.00E+07 | 690.3894 |
| **K11-GG** | TLTGKTITLEVEPSDTIENVKGG | ND | 1201.63668 |
| **K27-GG** | TITLEVEPSDTIENVKAKGG | ND | 1051.05479 |
| **K29-GG** | AKIQDKGG | ND | 408.73233 |
| **K33-GG** | IQDKEGIPPDQQRLIFAGKGG | ND | 1134.11077 |
| **K48-GG** | LIFAGKQLEDGRGG | 3.00E+08 | 730.89644 |
| **K63-GG** | TLSDYNIQKESTLHLVLRGG | ND | 1122.6027 |

**Supplementary Table 8.** Bottom-up analysis of ubiquitination sites for $Ub_n$-GFP mixture.

| | **Peptide** | **z** | **m/z** |
|---|---|---|---|
| **N term or K13** | GSSHHHHHHSMGKLGR | 3 | 639.63750 |
| **K180** | LEYNYNSHNVYIMADKQK | 3 | 782.37337 |
| **K262** | DHMVLLEFVTAAGITLGMDELYK | 3 | 894.45119 |
| **K186** | NGIKVNFK | 2 | 517.29309 |
| **K190** | VNFKIR | 1 | 890.52066 |
| **K125** | TIFFKDDGNYK | 2 | 731.35407 |
